# A non-canonical top-down pathway regulating relapse to opioid

**DOI:** 10.1101/2025.11.27.691060

**Authors:** Jing Liu, Ruo-Song Chen, Ru-Feng Ye, Gui-Ying Zan, Zi-Han Liu, Shuo Wu, Yi-Quan Shen, Xun-Qian Zhang, Chi Xu, Zhao-Qiu Wu, Yu-Jun Wang, Jing-Gen Liu

**Author notes:** These authors contributed equally (Jing Liu, Ruo-Song Chen). Corresponding author (Z.-Q.W.), (Y.-J.W.), (J.-G.L.).

## Abstract

The high relapse rates are major challenges in treating opioid addiction. However, it remains poorly understood the mechanisms underlying long-lasting vulnerability to relapse. Here, we identify the cortico-subthalamic projection from the granular retrosplenial cortex (RSG) to the zona incerta (ZI) as a key circuit for controlling relapse to opioid. We uncover that an ensemble of RSG glutamatergic neurons that locate in layer 5 and express B subtype cholecystokinin receptor (RSG^Glu-Cckbr^ ensemble) are critically implicated in the encoding and retrieval of opioid-associated memories, leading to relapse to opioid via innervating ZI GABAergic (ZI^GABA^) rather than dopaminergic neurons. We further identify that the Cckergic neurons from anterodorsal nucleus of thalamus (AD thalamus) orchestrates RSG^Glu-Cckbr^-ZI^GABA^ circuit function via release of cholecystokinin (Cck) in RSG. These results elucidate a previously unknown non-canonical top-down pathway regulating relapse to opioid and shed new light on the development of new therapeutic strategies to treat relapse to opioids.

## INTRODUCTION

Opioid addiction is a chronic, usually life-long disorder characterized by high rates of relapse to drug use despite successful abstinence^1^. A major challenge for treating drug addiction is to prevent relapse, which occurs largely due to the retrieval of long-lasting drug-associated memories that relate to drug-evoked experiences with environmental cues^2,3^. In humans, environments or contexts previously associated with drug use often provoke relapse after abstinence^4–7^. In rodents, exposure to the context paired with drug use after extinction reinstates drug seeking^8–10^. Drug-associated memories are encoded and retrieved by distributed neural networks that are not fully understood yet.

Compelling evidence suggests that addiction-related actions of abused drugs such as cocaine and opioids are controlled by interconnected prefrontal cortex (PFC) and mesolimbic brain regions, including the medial prefrontal cortex (mPFC), nucleus accumbens (NAc), ventral pallidum (VP) and ventral tegmental area (VTA)^11–15^. Although the reciprocal projections between mPFC, NAc and VTA are part of a canonical pathway that has been well established in the expression and reinstatement of drug-seeking behavior, accumulating evidence also suggests that dopamine– and mesocorticolimbic-independent substrates are involved in the addictive behaviors of opioids^16–20^. Nevertheless, little is currently known about their identity.

The retrosplenial cortex (RSC), a dorsomedial parietal area of the brain, including granular (RSG) and agranular (RSA) subregions, stands as a major hub for interconnectivity among multiple cortical and subcortical brain regions^21^. Anatomically, the RSC has widespread connections with the thalamus, subthalamus, hippocampus and cortices including visuo-sensory, postrhinal, anterior cingulate, and prefrontal cortex^22,23^. The RSC implements a variety of cognitive functions, including spatial navigation, encoding context value, storage and retrieval of contextual and episodic memory^24–28^. Although the RSC has also been reported to uniquely maintain value-related information in persistent population activity patterns and is required for reward-history-based decision making and behavioral strategy^29^, the specific role of RSC in neurocircuits responsible for encoding and retrieval of opioid-associated memories has not yet been studied.

In this study, we examined the role of the RSC in encoding and retrieval of contextual opioid-associated memories and found that a subset of RSG Cckbr-positive glutamatergic neurons underlies the encoding and retrieval of opioid-associated memories via targeting the zona incerta (ZI) GABAergic rather than dopaminergic neurons in the rat brain. The activity of RSG-ZI circuit is orchestrated by cholecystokinin (Cck) derived from the anterodorsal nucleus of thalamus (AD thalamus). Our results establish RSG as a critical node in the neural networks involved in the encoding, storage and retrieval of opioid-associated memories through its afferent connection with the AD thalamus and efferent connection with the ZI.

## RESULTS

### RSG^Glu^ neurons are required for the encoding and retrieval of opioid-associated memories

Given that RSC uniquely maintains persistent value-related population activity and is required for reward-history-based decision making and behavioral strategy^29^, we hypothesized that RSC might be involved in the encoding and retrieval of opioid-associated memories. To test this, we first determined the effects of heroin self-administration (heroin SA) training and re-exposure of rats that underwent heroin SA training to context paired with opioid use after extinction (context-induced heroin relapse, CIR) on the neuronal activity of RSC by examination of c-fos expression with immunostaining and by recording of Ca^2+^ signal with in vivo fiber photometry. To do this, two different groups of animals were used for the experiments: Heroin-ABB and Heroin-ABA. In the Heroin-ABB group, rats were trained to self-administer heroin in context A and extinguished in context B. In the Heroin-ABA group, rats underwent heroin SA training in context A, extinction in context B and ignition of craving for heroin in context A. Rats of both groups were subjected to c-fos immunostaining (Figure S1A). A significantly increased number of c-fos positive (c-fos^+^) cells was observed in RSG but not RSA of CIR rats (Figures S1B-S1D), suggesting that CIR induced the remarkable excitation of RSG rather than RSA. We then tested the co-localization of c-fos with CaMKIIα, a specific marker of glutamatergic neurons, and found that >90% of c-fos^+^ cells were co-localized with CaMKIIα (Figure S1E), indicating the activation of RSG glutamatergic neurons (RSG^Glu^). Using fluorescent *in situ* hybridization (FISH), we further found that CIR resulted in robust activation of Vglut1-positive glutamatergic (RSG^Vglut1^) neurons instead of Vgat-positive GABAergic neurons in RSG (Figures 1A and 1B). These results suggest that RSG^Vglut1^ neurons are probably involved in encoding and retrieval of heroin-associated memories.

**Figure 1.**
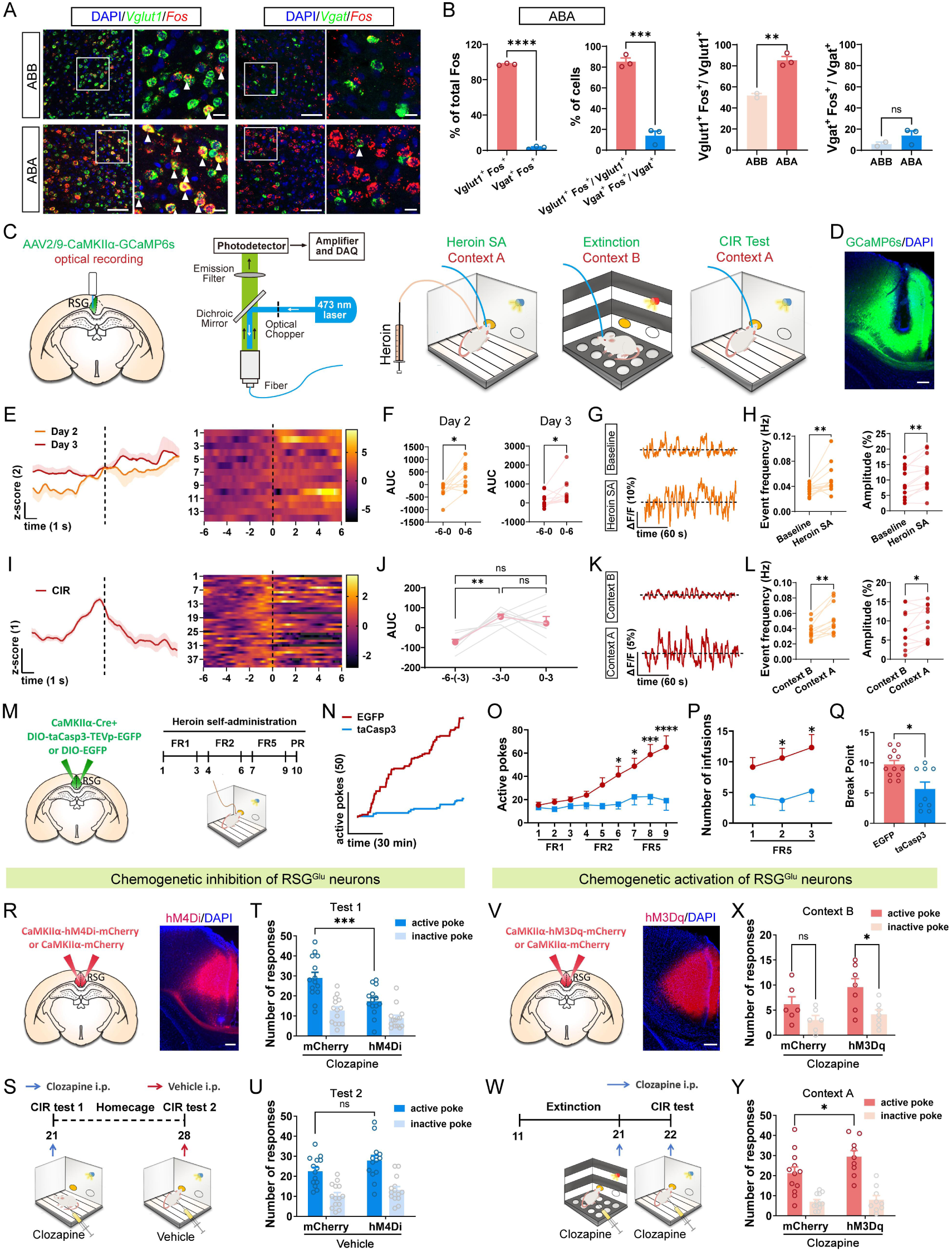
RSG glutamatergic neurons are critically required for the encoding and retrieval of contextual heroin memory. **(A)** Representative images showing the co-localization of *Fos* with *Vglut1* or *Vgat* in RSG of ABB and ABA groups. Scale bars, 100 μm and 20 μm. **(B)** Percentage of Vglut1^+^ Fos^+^ and Vgat^+^ Fos^+^ neurons in ABA (*n* = 3) and ABB (*n* = 2) groups. Unpaired t test, \*\**p* < 0.01, \*\*\**p* < 0.001, \*\*\*\**p* < 0.0001, ns, no significant difference. **(C)** Schematic showing the viral strategy, fiber implantation and schedule of fiber photometry recording. **(D)** Representative image showing the expression of GCaMP6s in RSG. Scale bars, 200 μm. **(E-F)** Average Ca^2+^ signals, heatmaps (E) and area under curve (F) of RSG glutamatergic neurons during day 2 (*n* = 10) and day 3 (*n* = 11) heroin SA training sessions. Vertical dashed line indicates the onset of active poke behaviors. Wilcoxon matched-pairs signed rank test, \**p* < 0.05. **(G-H)** Representative traces (G) and the event frequency and average amplitude (H) of the tonic Ca^2+^ signals during baseline recording and day 3 of heroin SA training (*n* = 12). For event frequency, Wilcoxon matched-pairs signed rank test, \*\**p* < 0.01. For amplitude, Paired t test, \*\**p* < 0.01. **(I-J)** Average Ca^2+^ signals, heatmaps (I) and area under curve (J) of RSG glutamatergic neurons during context-induced relapse test (*n* = 8). Vertical dashed line indicates the onset of active poke behaviors. RM One-way ANOVA (F_(1.418, 9.923)_ = 7.671, *p* < 0.05) followed by Tukey’s post hoc test, \*\**p* < 0.01, ns, no significant difference. **(K-L)** Representative traces (K) and the event frequency and average amplitude (L) of the tonic Ca^2+^ signals during extinction training in context B and relapse test in context A (*n* = 11). For event frequency, Wilcoxon matched-pairs signed rank test, \*\**p* < 0.01. For amplitude, Paired t test, \**p* < 0.05. **(M)** Schematic showing the viral strategy for selective ablation of RSG glutamatergic neurons and the training schedule for heroin SA. **(N)** Representative cumulative curve of the number of active poke behaviors during FR5 heroin SA training. **(O)** Number of active poke behaviors in EGFP control group (*n* = 13) and taCasp3 group (*n* = 10) during heroin SA training under FR1-FR2-FR5 schedule. Two-way ANOVA (F_(1,188)_ = 50.61, *p* < 0.0001) followed by Sidak’s post hoc test, \**p* < 0.05, \*\*\**p* < 0.001, \*\*\*\**p* < 0.0001. **(P)** Number of heroin infusions in EGFP control group (*n* = 13) and taCasp3 group (*n* = 10) during heroin SA training under FR5 schedule. Two-way ANOVA (F_(1,62)_ = 20.22, *p* < 0.0001) followed by Sidak’s post hoc test, \**p* < 0.05. **(Q)** Number of break point in EGFP control group (*n* = 12) and taCasp3 group (*n* = 9) during heroin SA training under PR schedule. Mann-Whitney test, \**p* < 0.05. **(R)** Left: schematic showing the viral strategy for chemogenetic inhibition of RSG glutamatergic neurons. Right: representative image showing the expression of hM4Di in RSG. Scale bars, 200 μm. **(S)** The test schedule for context-induced relapse. **(T)** Number of responses in mCherry control group (*n* = 14) and hM4Di group (*n* = 14) during test 1 with clozapine injection (i.p.). Two-way ANOVA (F(_1,52)_ = 14.51, *p* < 0.001) followed by Sidak’s post hoc test, \*\*\**p* < 0.001. **(U)** Number of responses in mCherry control group (*n* = 14) and hM4Di group (*n* = 14) during test 2 with vehicle injection (i.p.). Two-way ANOVA (F_(1,52)_ = 4.297, *p* = 0.0431) followed by Sidak’s post hoc test, ns, no significant difference. **(V)** Left: schematic showing the viral strategy for chemogenetic activation of RSG glutamatergic neurons. Right: representative image showing the expression of hM3Dq in RSG. Scale bars, 200 μm. **(W)** The test schedule in context B or context A. **(X)** Number of responses of mCherry control group (*n* = 6) and hM3Dq group (*n* = 7) in context B with clozapine injection (i.p.). Two-way RM ANOVA (F_(1,12)_ = 12.84, *p* < 0.01) followed by Sidak’s post hoc test, \**p* < 0.05. **(Y)** Number of responses of mCherry control group (*n* = 12) and hM3Dq group (*n* = 9) in context A with clozapine injection (i.p.). Two-way ANOVA (F_(1,38)_ = 3.444, *p* = 0.0712) followed by Sidak’s post hoc test, \**p* < 0.05.

Next, we employed fiber photometry to record Ca^2+^ signals in vivo to further validate the activation of RSG^Glu^ neurons by heroin SA training and CIR. Adeno-associated virus (AAVs) carrying the genetically encoded Ca^2+^ indicator (GCaMP6s) with CaMKIIα promoter were injected into RSG or RSA followed by optical fibers implantation (Figures 1C, 1D, S2A and S2B). The specificity of the virus carrying CaMKIIα promoter was verified by FISH (Figures S1F and S1G). We found that Ca^2+^ signals in RSG but not RSA were significantly increased both during the early period of heroin SA training (day 2-3) (Figures 1E-1H, S2C and S2D) and during CIR (Figures 1I-1L, S2E and S2F). Significant increase of RSG Ca^2+^ signals were accompanied by conducting active nose pokes, suggesting that RSG^Glu^ but not RSA^Glu^ neurons participate in the encoding and retrieval of heroin-associated memories. We also found that heroin SA training and CIR had no significant effect on the Ca^2+^ activity of GABAergic neurons in RSG (RSG^GABA^ neurons), measured by fiber photometry recording via delivering AAV-Vgat-GCaMP6s into RSG (Figures S2G-S2O).

Next, we investigated the role of RSG^Glu^ neurons in the encoding and retrieval of heroin-associated memories by testing heroin SA and CIR behaviors. We first examined the role of RSG^Glu^ neurons in heroin SA behaviors by employing the Cre-dependent taCasp3-TEVp system to selectively ablate RSG^Glu^ neurons through bilateral injections of AAV-CaMKIIα-Cre and AAV-DIO-taCasp3-TEVp-EGFP or AAV-DIO-EGFP into RSG (Figure 1M). The efficiency of taCasp3-TEVp system was verified by immunostaining of CaMKIIα in RSG, which showed that the number of CaMKIIα-positive neurons was robustly decreased in the taCasp3 group compared to the EGFP control group, indicating the ablation of glutamatergic neurons in RSG (Figures S3A and S3B). We then trained the rats to self-administer heroin under FR1-FR2-FR5-PR schedule (Figure 1M). Although the heroin taking under FR1 was not affected, ablation of RSG^Glu^ neurons dramatically suppressed the active poke behaviors after the schedule was shifted to FR2 and FR5 (Figures 1N and 1O). The number of heroin infusions under FR5 and the break point under PR program were also significantly decreased (Figures 1P and 1Q). These results indicate that RSG^Glu^ neurons are essential for the encoding of heroin SA behavior.

Next, we determined whether RSG^Glu^ neurons are also required for retrieval of heroin-associated memories by examining CIR behaviors with chemogenetic manipulations of RSG^Glu^ neurons. To do this, we injected AAV-CaMKIIα-hM4Di-mCherry, AAV-CaMKIIα-hM3Dq-mCherry or AAV-CaMKIIα-mCherry into the RSG to silence or activate RSG^Glu^ neurons. We found that chemogenetically inhibiting RSG^Glu^ neurons significantly reduced the active poke of hM4Di-expressing rats compared to the mCherry controls (test 1) without affecting the locomotion (Figures 1R-1T, S3C and S3D). To determine whether heroin-associated memories of rats could be retrieved when the RSG^Glu^ neurons were not inhibited, these rats were kept in their homecages for 7 days and then they were subjected to the context paired with heroin taking to retrieve memories again under the condition without clozapine injection (test 2). As expected, the heroin seeking behaviors were recovered (Figure 1U), indicating that heroin-associated memories encoded by RSG^Glu^ neurons are necessary to relapse to heroin seeking. The importance of RSG^Glu^ neurons for the retrieval of heroin-associated memories was further supported by the fact that chemogenetical activation of RSG^Glu^ neurons also induced relapse behaviors in context B after the drug seeking behaviors were extinguished in this environment and that activation of RSG^Glu^ neurons further increased active poke behaviors as measured in context A (Figures 1V-1Y) without affecting the locomotion of rats (Figures S3E and S3F). Functional validation of hM3Dq in RSG^Glu^ neurons was confirmed by c-fos immunostaining (Figures S3G and S3H). Together, these results demonstrate that RSG^Glu^ neurons are required for the encoding of heroin SA behavior and context-induced relapse.

### The transcriptionally specific ensemble of RSG^Vglut1^ neurons was recruited by context-induced relapse to heroin

Compelling evidence has suggested the cellular and transcriptional heterogeneity of the cortical neurons^20^. Genetically distinct populations of neurons in cortex are linked to diverse physiological and pathological functions^20,30,31^. To identify genetically specific RSG glutamatergic ensembles that are critically implicated in the encoding and retrieval of heroin-associated memories, leading to relapse, we began by utilizing single-nuclei RNA sequencing (snRNA-seq) to characterize the transcriptional profiles of these cells. We micro-dissected and collected the RSG tissues from brain slices of adult rats (Figure S4A), and then isolated the single nuclei to obtain information of transcriptomes and generate cDNA libraries for subsequent analysis (Figure 2A). Following computational removing of low-quality cells and doublets, 35019 single cells were achieved and identified to seven discrete clusters in uniform manifold approximation and projection (UMAP) space based on the expression of typical cell markers including: neuron (*Snap25*), astrocyte (*Gfap*), microglia (*Dock2*), oligodendrocyte (*Mog*), oligodendrocyte precursor cell (*Pdgfra*), endothelial cell (*Prom1*), fibroblast (*Col1a1*) (Figures S4B-S4D).

**Figure 2.**
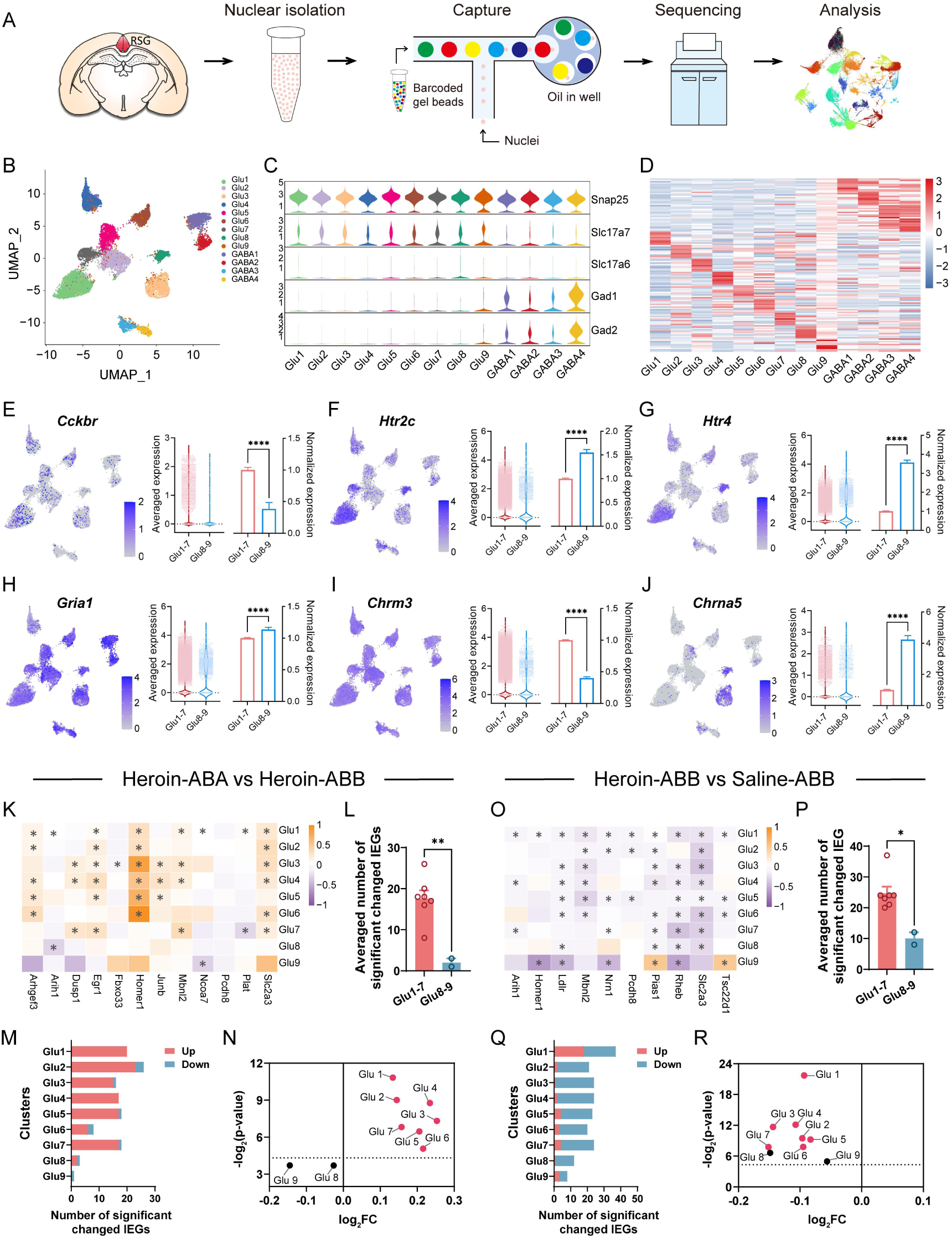
Context-induced heroin relapse recruits transcriptionally specific glutamatergic neuronal ensemble in RSG. **(A)** Schematic showing the workflow for snRNA-seq. **(B)** Transcriptional profiles of RSG neurons visualized by Uniform manifold approximation and projection (UMAP) dimensional reduction. **(C)** Violin plots showing the expression pattern of marker genes for glutamatergic neurons and GABAergic neurons in RSG. **(D)** Heatmaps showing the expression of top ten marker genes in different neuronal clusters. **(E-J)** Feature plots showing the expression patterns of *Cckbr* (E), *Htr2c* (F), *Htr4* (G), *Gria1* (H), *Chrm3* (I) *and Chrna5* (J) in each neuronal cluster and the expression differences between clusters Glu 1-7 *vs* Glu 8-9. Mann-Whitney test, \*\*\*\**p* < 0.0001. **(K, O)** Heatmaps showing the expression difference of the representative IEGs [Fold Change (FC) > 1.2 in at least one cluster] between Heroin-ABA group *vs* Heroin-ABB group (K) or Heroin-ABB group *vs* Saline-ABB group (O). Wilcoxon rank-sum test followed by Benjamini-Hochberg post hoc test, \**p* < 0.05. **(L, P)** Number of significantly changed IEGs in different clusters of RSG neurons compared between Heroin-ABA group *vs* Heroin-ABB group (L) or Heroin-ABB group *vs* Saline-ABB group (P). One circle indicates an individual glutamatergic neuronal cluster. Unpaired t test, \**p* < 0.05, \*\**p* < 0.01. **(M, Q)** Number of significantly up– or down-regulated IEGs in each glutamatergic neuronal cluster compared between Heroin-ABA group *vs* Heroin-ABB group (M) or Heroin-ABB group *vs* Saline-ABB group (Q). **(N, R)** Average log_2_FC and –log_2_(*p*-value) of the top-10 differentially expressed IEGs in each glutamatergic neuronal cluster compared between Heroin-ABA group *vs* Heroin-ABB group (N) or Heroin-ABB group *vs* Saline-ABB group (R).

To characterize the transcriptional profiles of these neuronal cell types in detail, we re-clustered the neuronal populations based on gene expression. Nuclei from a total of 22,148 neuronal cells were included in the analyses after filtering for quality control. Cells were segregated into 13 discrete clusters in UMAP space according to their transcriptional profiles (Figures 2B and S4E-S4I). Gene transcripts considered hallmarks of glutamatergic neurons were concentrated in clusters 1-9 and those of GABAergic neurons were concentrated in clusters 10-13 (Figure 2C). Glutamatergic neurons in clusters 1-9 (Glu 1-9) predominantly expressed *Slc17a7* that encodes vesicular glutamate transporter 1 (*Vglut1*). Cells in clusters 10-13 (GABA 1-4) expressed both glutamate decarboxylase 1 (*Gad1*) or glutamate decarboxylase 2 (*Gad2*), the markers of GABAergic neurons (Figure S4J). Conserved marker analysis was subsequently performed to define gene signatures characterizing each cluster (Figures 2D and S4K-S4M). Several gene transcripts, which encode the B subtype of cholecystokinin receptor (*Cckbr*, *Cckar* were absent in RSG), Serotonin (5-Hydroxytryptamine) receptor 2c and 4 (*Htr2c*, *Htr4*), Glutamate ionotropic receptor AMPA type subunit 1 (*Gria1*), Cholinergic receptor muscarinic 3 and nicotinic alpha 5 (*Chrm3*, *Chrna5*), were also detected in clusters Glu 1-9 (Figures 2E-2J). Interestingly, distinctly expressing profiles were observed between clusters Glu 1-7 and Glu 8-9, with a great amount of *Cckbr* and *Chrm3* expressed in clusters Glu 1-7 and higher expression of *Htr2c*, *Htr4, Gria1, Chrna5* in clusters Glu 8-9 (Figures 2E-2J). Additionally, by use of FISH, we verified that the Cckbr-positive cells in RSG were predominantly Vglut1-positive glutamatergic neurons (Figure S5A), with a small amount of Cckbr-positive cells being GABAergic neurons (Figure S5B). More importantly, we also found that heroin SA training and CIR significantly increased the average *Cckbr* mRNA in clusters Glu 1-7 rather than Glu 8-9, compared to the saline SA training group (Figures S5C and S5D). Correspondingly, the elevation of Cckbr protein expression is also detected by immunostaining of Cckbr in RSG (Figures S5E and S5F). These results suggest that heroin SA training and CIR might exert an effect on RSG^Vglut1^ neurons through regulation of Cckbr expression.

Next, we detected the effects of CIR on the neuronal activity of the clusters Glu 1-7 and clusters Glu 8-9 neurons by examining the expression of the immediate early genes (IEGs). To do this, three experimental groups were used: Saline-ABB, Heroin-ABB and Heroin-ABA. In Saline-ABB group, rats were trained to self-administer saline in context A, and then extinguished in context B. All groups were confirmed equal representation of neuronal clusters, proportion of neurons and gene and UMI counts in each neuronal cluster (Figures S6A-S6D). Patterns and relative densities of clusters were similar between Heroin-ABA, Heroin-ABB and Saline-ABB groups (Figure S6B). This indicates that the extinction of heroin SA behavior and the CIR test didn’t change the overall neuronal types and transcriptional features of RSG.

We then systematically profiled and compared the expression of the immediate early genes (IEGs), markers of neuronal activation, in each neuronal cluster between the three groups. While the *Fos* expression detected was low (1.5% of neurons), possibly due to the *Fos* depletion in the nucleus, we observed several alternative IEGs with relatively higher expression, such as: *Arc* (2.0%), *Egr1* (9.8%), and *Homer1* (51.9%) (Figure S6E). We therefore expanded our analysis according to the IEGs inventory reported previously^32,33^. First, we evaluated changes in IEGs expression induced by CIR via comparing alteration of IEGs expression between Heroin-ABA and Heroin-ABB groups in each cluster of neurons. CIR resulted in upregulation of a great number of IEGs in clusters Glu 1-7 relative to clusters Glu 8-9 (Figures 2K-2M and Table 1). Analysis of the averaged fold change and *p*-value of the top 10 differentially expressed IEGs in each glutamatergic cluster revealed that each cluster in Glu 1-7 in Heroin-ABA group displayed obviously increased average IEG expression compared to Heroin-ABB group, except for clusters Glu 8-9 (Figure 2N). These results indicate that CIR selectively increases neuronal activity of the clusters Glu 1-7. Next, to investigate whether the clusters Glu 1-7 neurons were also affected by extinction of heroin SA behavior, we compared the change of IEGs expression occurred in Heroin-ABB and Saline-ABB groups and found that like CIR, extinction training also altered IEGs expression in clusters Glu 1-7, indicated by a remarkably decreased average IEGs expression compared to Saline-ABB group (Figures 2O-2R and Table 2). Moreover, employing Kyoto Encyclopedia of Genes and Genomes (KEGG) enrichment analysis, we found that the CIR-induced differential expressed genes occurred in clusters Glu 1-7 compared to Glu 8-9 were enriched in the Morphine Addiction pathway (Figures S6F and S6G). Taken together, these results demonstrated that RSG^Vglut1^ neurons are transcriptional heterogeneity and suggested that RSG Cckbr-positive glutamatergic (RSG^Glu-Cckbr^) neurons may be potentially implicated in the encoding and retrieval of heroin-associated memory.

### RSG^Glu-Cckbr^ neurons may critically contribute to the encoding, storage and retrieval of opioid-associated memory

To determine whether RSG^Glu-Cckbr^ neurons contribute to the formation of heroin SA behavior and CIR, we employed the gene-encoded neurotransmitter sensors combined with fiber photometry to monitor the release of Cck, 5-HT, glutamate and acetylcholine, the endogenous ligands of cholecystokinin receptor, serotonin receptor, AMPA receptor and cholinergic receptor (Figures 3A-3C). We found that the signal of Cck rather than 5-HT, glutamate and acetylcholine were robustly increased in response to the active poke behaviors during both the early formation period of heroin SA (day 2-3) and CIR (Figures 3D-3K), indicating the selective increase of Cck concentration in RSG during encoding and retrieval of heroin-associated memories. We next tested neuronal activity of RSG^Glu-Cckbr^ neurons following heroin SA training and CIR by employing FISH and in vivo fiber photometry recording. We found that, corresponding to the enhancement of Cck concentrations, the activities of RSG^Glu-Cckbr^, but not RSG Cckbr-negative glutamatergic neurons were increased significantly, manifested by a remarkable increase in Ca^2+^ signal of RSG^Glu-Cckbr^ neurons during both the early formation period of heroin SA and CIR (Figures 3L-3U and S7A-S7J). Activation of RSG^Glu-Cckbr^ neurons by CIR was further confirmed by increased *Fos* expression in the RSG^Glu-Cckbr^ neurons (Figure 3V). These results suggest the activation of RSG^Glu-Cckbr^ neurons may be required for encoding and retrieval of heroin-associated memories. Interestingly, activation of the RSG^Glu-Cckbr^ neurons seems not to be required for natural rewards, as these neurons were inhibited by water licking and chocolate consumption (Figures S8A-S8D).

**Figure 3.**
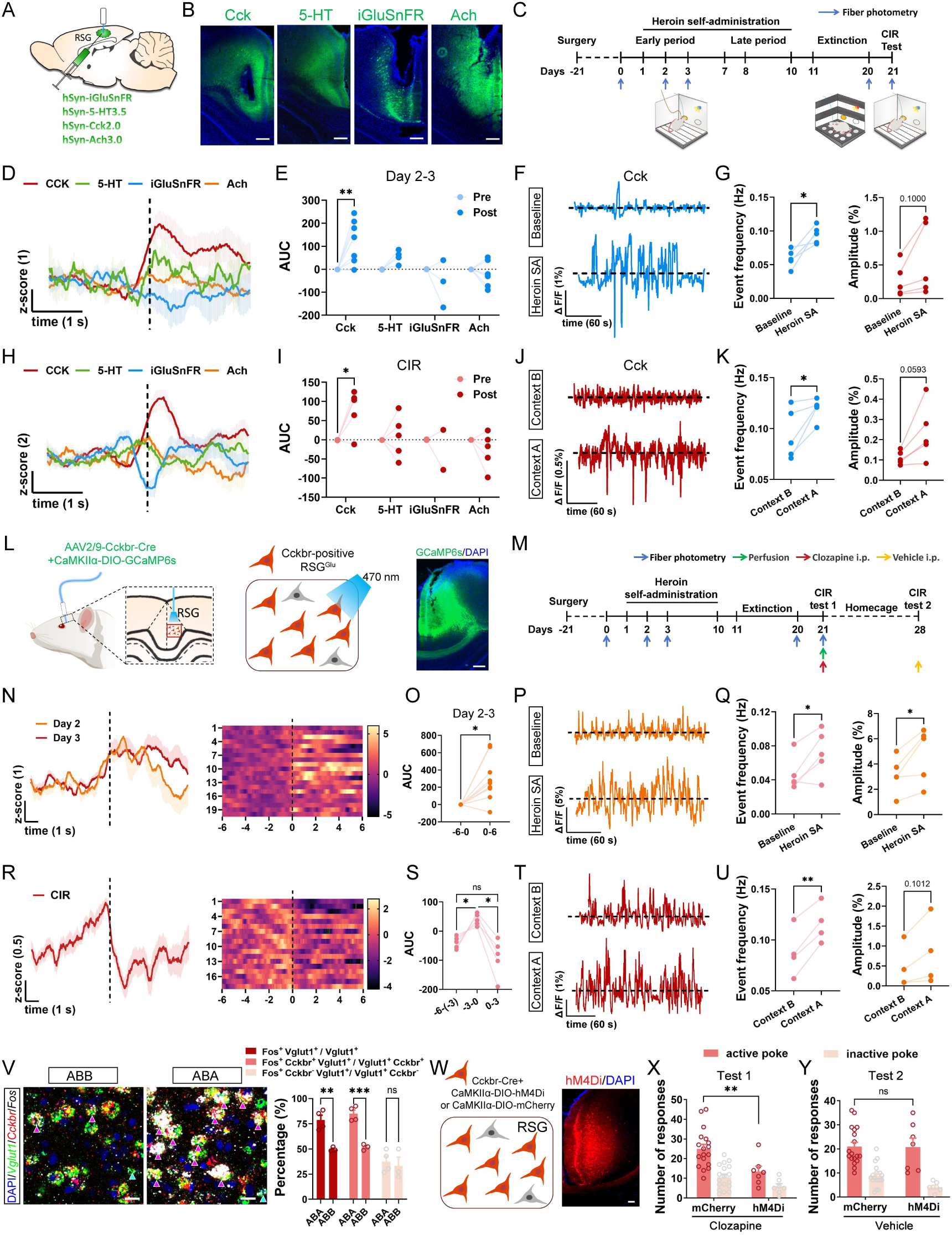
The Cckbr-positive glutamatergic neuronal ensemble in RSG specifically contributes to the encoding and retrieval of contextual heroin memory. **(A)** Schematic showing the viral strategy and fiber implantation for fiber photometry recording of different neurotransmitters in RSG. **(B)** Representative images showing the expression of different neurotransmitter sensors in RSG. Scale bars, 200 μm. **(C)** Schematic showing the recording schedule during heroin SA training, extinction and context-induced relapse test. **(D-E)** Average signals (D) and area under curve (E) of the Cck (*n* = 8), 5-HT (*n* = 4), glutamate (*n* = 3) and acetylcholine (*n* = 6) during early (day 2-3) phase of heroin SA training. Two-way RM ANOVA (F_(3,17)_ = 4.746, *p* < 0.05) followed by Sidak’s post hoc test, \*\**p* < 0.01. Vertical dashed line indicates the onset of active poke behaviors. **(F-G)** Representative traces (F) and the event frequency and average amplitude (G) of the tonic Cck signals during baseline recording and day 3 of heroin SA training (*n* = 5). Paired t test, \**p* < 0.05. **(H-I)** Average signals (H) and area under curve (I) of the Cck (*n* = 5), 5-HT (*n* = 5), glutamate (*n* = 2) and acetylcholine (*n* = 5) during context-induced relapse. Two-way RM ANOVA (F_(3,13)_ = 3.689, *p* < 0.05) followed by Sidak’s post hoc test, \**p* < 0.05. Vertical dashed line indicates the onset of active poke behaviors. **(J-K)** Representative traces (J) and the event frequency and average amplitude (K) of the tonic Cck signals during extinction training and relapse test (*n* = 5). Paired t test, \**p* < 0.05. **(L)** Left: schematic showing the viral strategy and fiber implantation for Ca^2+^ signal recording of Cckbr-positive glutamatergic neurons in RSG. Right: representative image showing the expression of GCaMP6s in RSG. Scale bars, 200 μm. **(M)** Schematic showing the schedule for fiber photometry recording, behavioral training and perfusion. **(N-O)** Average Ca^2+^ signals, heatmaps (N) and area under curve (O) of RSG Cckbr-positive glutamatergic neurons during day 2-3 of heroin SA training (*n* = 10). Vertical dashed line indicates the onset of active poke behaviors. Paired t test, \**p* < 0.05. **(P-Q)** Representative traces (P) and the event frequency and average amplitude (Q) of the tonic Ca^2+^ signals during baseline recording and day 3 of heroin SA training (*n* = 5). Paired t test, \**p* < 0.05. **(R-S)** Average Ca^2+^ signals, heatmaps (R) and area under curve (S) of RSG Cckbr-positive glutamatergic neurons during context-induced relapse test (*n* = 5). Vertical dashed line indicates the onset of active poke behaviors. RM One-way ANOVA (F_(1.183, 4.734)_ = 11.96, *p* < 0.05) followed by Tukey’s post hoc test, \**p* < 0.05, ns, no significant difference. **(T-U)** Representative traces (T) and the event frequency and average amplitude (U) of the tonic Ca^2+^ signals of RSG Cckbr-positive glutamatergic neurons during extinction training and relapse test (*n* = 4). Paired t test, \*\**p* < 0.01. **(V)** Left: representative images showing the expression of *Fos*, *Vglut1* and *Cckbr* in RSG in ABB and ABA groups. The arrows indicate Vglut1^+^ Cckbr^+^ Fos^+^ cells (pink) and Vglut1^+^ Cckbr^-^ Fos^+^ (green) neurons. Scale bars, 20 μm. Right: percentage of Fos^+^ cells in Vglut1^+^, Vglut1^+^ Cckbr^+^ and Vglut1^+^ Cckbr^-^ neurons in RSG of ABB (*n* = 3) *vs* ABA (*n* = 4) groups. Two-way ANOVA (F_(1,15)_ = 30.55, *p* < 0.0001) followed by Sidak’s post hoc test, \*\**p* < 0.01, \*\*\**p* < 0.001, ns, no significant difference. **(W)** Left: schematic showing the viral strategy for selectively inhibition of RSG Cckbr-positive glutamatergic neurons. Right: representative image showing the expression of hM4Di in RSG. Scale bars, 200 μm. **(X)** Number of responses in mCherry control group (*n* = 18) and hM4Di group (*n* = 7) during test 1 with clozapine injection (i.p.). Two-way ANOVA (F_(1,46)_ = 11.69, *p* < 0.01) followed by Sidak’s post hoc test, \*\**p* < 0.01. **(Y)** Number of responses in mCherry control group (*n* = 18) *vs* hM4Di group (*n* = 7) during test 2 with vehicle injection (i.p.). Two-way ANOVA (F_(1,46)_ = 1.433, *p* = 0.2374) followed by Sidak’s post hoc test, ns, no significant difference.

Next, we verified the role of RSG^Glu-Cckbr^ neurons in the retrieval of heroin-associated memory by examining the effect of chemogenetic inhibition of RSG^Glu-Cckbr^ neurons on the CIR behavior (Figure 3W). As expected, inhibition of RSG^Glu-Cckbr^ neurons significantly decreased the number of active pokes during CIR tests (Figure 3X). However, heroin-associated memory could be retrieved again 7 days later when the RSG^Glu-Cckbr^ neurons were not inhibited with vehicle injection (test 2) (Figure 3Y), suggesting that heroin-associated memories may be stored in RSG^Glu-Cckbr^ neurons. Although there is also a small proportion (29%) of RSG GABAergic neurons (RSG^GABA-Cckbr^) that also express low levels of *Cckbr* (Figure S5B), in vivo fiber photometry recording demonstrated that heroin SA training failed to activate RSG^GABA-Cckbr^ neurons (Figures S9A-S9G) and chemogenetic inhibition of RSG^GABA-Cckbr^ neurons had no significant effect on the retrieval of heroin-associated memories (Figures S9H-S9K). Overall, these results suggest that RSG^Glu-Cckbr^ neurons may critically contribute to the encoding, storage and retrieval of heroin-associated memory.

### CRISPR-Cas9-mediated knockdown of Cckbr in RSG^Glu^ neurons suppressed context-induced relapse to heroin seeking

Cckbr is widely distributed in cortical and subcortical brain regions and critically involved in long-term potentiation and memory formation^34,35^. In addition, Cck has also been shown to be involved in regulation of opioid addictive behaviors^36^. To further confirm the importance of Cck/Cckbr signaling for the role of RSG^Glu-Cckbr^ neurons in encoding and retrieval of heroin-associated memories, we examined the effect of pharmacological and genetical manipulations of Cck/Cckbr signaling on heroin SA and CIR-induced changes in the activity of RSG^Glu-Cckbr^ neurons and CIR behaviors. We first microinfused L365260, a selective Cckbr antagonist, into RSG and found that blockade of Cck/Cckbr signaling significantly decreased the relapse behavior (Figures S10A-S10C). Next, we knocked down Cckbr in the RSG^Glu^ neurons by delivering AAVs carrying shRNA (scramble-shRNA or Cckbr-shRNA) with CaMKIIα promoter into RSG (Figure 4A). The knockdown efficiency was verified by FISH (Figures 4B and 4C). We trained the rats under FR1-FR2-PR schedule 5 weeks after AAVs delivery or under FR1 schedule 2 weeks after AAVs delivery followed by extinction training and CIR test (Figures 4D and 4F). We found that Cckbr-shRNA-infected rats showed a significant reduction of both heroin SA behavior and context-induced relapse to heroin, compared with that in rats expressing a control scrambled shRNA (Figures 4E and 4G). Correspondingly, knockdown of Cckbr with Cckbr-shRNA also abolished heroin SA training and CIR-induced increase in Ca^2+^ signal during the early formation period of heroin SA and during CIR (Figures 4H-4Q).

**Figure 4.**
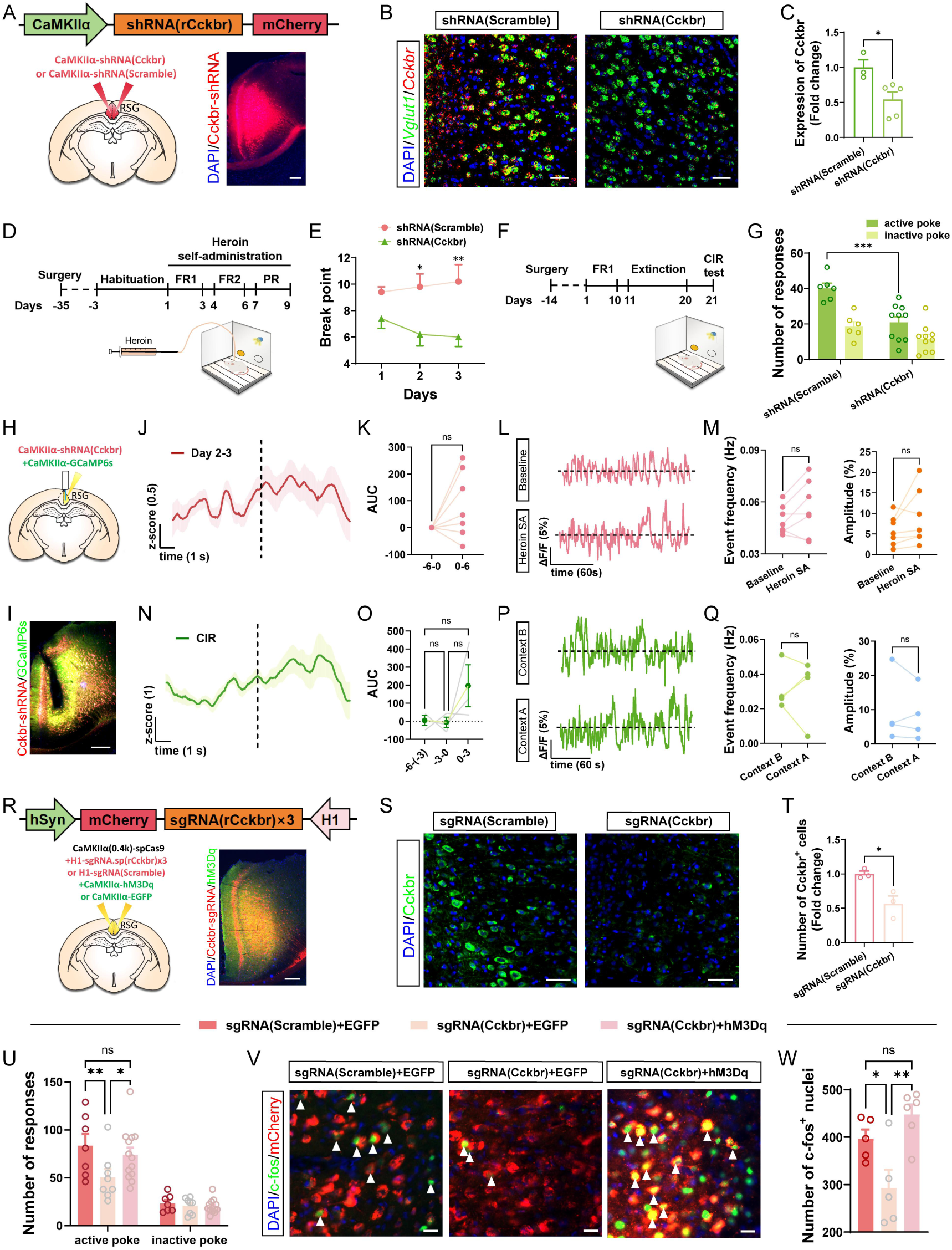
Cckbr in RSG modulates contextual heroin memory retrieval via regulating the activity of glutamatergic neurons. **(A)** Schematic showing the viral strategy for *Cckbr* knockdown in RSG glutamatergic neurons and the representative image of the virus expression in RSG. Scale bars, 200 μm. **(B)** Representative images of *Cckbr* mRNA expression in RSG Vglut1^+^ neurons in shRNA(scramble)– and shRNA(Cckbr)-expressing rats. Scale bars, 50 μm. **(C)** Quantification of *Cckbr* mRNA in RSG Vglut1^+^ neurons of shRNA(scramble) (*n* = 3) and shRNA(Cckbr) (*n* = 5) expressing rats. Unpaired t test, \**p* < 0.05. **(D)** Schematic showing the training schedule of heroin SA under FR1-FR2-PR program. **(E)** Break points of shRNA(Scramble) (*n* = 5) and shRNA(Cckbr) (*n* = 5) expressing rats during 3-day PR training sessions. Two-way ANOVA (F_(1,24)_ = 21.15, *p* < 0.001) followed by Sidak’s post hoc test, \**p* < 0.05, \*\**p* < 0.01. **(F)** Schematic showing the training schedule of heroin SA under FR1 program, extinction in context B and context-induced relapse in context A. **(G)** Number of responses of shRNA(Scramble) (*n* = 6) and shRNA(Cckbr) (*n* = 10) expressing rats during context-induced relapse test. Two-way ANOVA (F_(1,28)_ = 18.36, *p* < 0.001) followed by Sidak’s post hoc test, \*\*\**p* < 0.001. **(H)** Schematic showing the viral strategy and fiber implantation in RSG for *Cckbr* knockdown combined with Ca^2+^ recording. **(I)** Representative image showing the expression of GCaMP6s and shRNA(Cckbr) in RSG. Scale bars, 200 μm. **(J-K)** Average Ca^2+^ signal (J) and area under curve (K) of the RSG glutamatergic neurons with *Cckbr* knockdown during day 2-3 of heroin SA training (*n* = 7). Paired t test, ns, no significant difference.Vertical dashed line indicates the onset of active poke behaviors. **(L-M)** Representative traces (L) and the event frequency and average amplitude (M) of the tonic Ca^2+^ signals of the RSG glutamatergic neurons with *Cckbr* knockdown during baseline recording *vs* heroin SA training (*n* = 7). Paired t test, ns, no significant difference. **(N-O)** Average Ca^2+^ signal (N) and area under curve (O) of the RSG glutamatergic neurons with *Cckbr* knockdown during context-induced relapse (*n* = 3). RM One-way ANOVA (F_(1.403, 2.087)_ = 2.435, *p* = 0.2562) followed by Tukey’s post hoc test, ns, no significant difference. Vertical dashed line indicates the onset of active poke behaviors. **(P-Q)** Representative traces (P) and the event frequency and average amplitude (Q) of the tonic Ca^2+^ signals of the RSG glutamatergic neurons with *Cckbr* knockdown during extinction training *vs* relapse test (*n* = 4). Paired t test, ns, no significant difference. **(R)** Schematic showing the viral strategy for CRISPR-Cas9-mediated knockdown of *Cckbr* and the chemogenetic activation of RSG glutamatergic neurons and the representative image of virus expression in RSG. Scale bars, 200 μm. **(S)** Representative images of Cckbr expression in RSG of sgRNA(Scramble)– and sgRNA(Cckbr)-expressing rats. Scale bars, 50 μm. **(T)** Quantification of Cckbr protein in RSG of sgRNA(Scramble) (*n* = 3) and sgRNA(Cckbr) (*n* = 3) expressing rats. Unpaired t test, \**p* < 0.05. **(U)** Number of responses in rats of control group (*n* = 7), Cckbr knockdown group (*n* = 8) and Cckbr knockdown with hM3Dq group (*n* = 13) during context-induced relapse. Two-way ANOVA (F_(2,50)_ = 3.033, *p* = 0.0571) followed by Sidak’s post hoc test, \**p* < 0.05, \*\**p* < 0.01, ns, no significant difference. **(V)** Representative images showing the c-fos expression in rats of control group, Cckbr knockdown group and Cckbr knockdown with hM3Dq group during context-induced relapse. Scale bars, 20 μm. **(W)** Number of c-fos positive nuclei in control group (*n* = 5), Cckbr knockdown group (*n* = 5) and Cckbr knockdown with hM3Dq group (*n* = 6). One-way ANOVA (F_(2, 13)_ = 8.678, *p* < 0.01) followed by Tukey’s post hoc test, \**p* < 0.05, \*\**p* < 0.01, ns, no significant difference.

To further confirm the pivotal role of Cck/Cckbr signaling in regulation of activity of RSG^Glu^ neurons and CIR behaviors, we employed CRISPR-Cas9-mediated gene editing combined with chemogenetic manipulation (Figure 4R). Knockdown efficiency of Cckbr by CRISPR-Cas9 was verified by immunostaining in RSG (Figures 4S and 4T). We found that the CRISPR-Cas9-mediated knockdown of Cckbr in RSG^Glu^ neurons significantly inhibited context-induced activation of RSG^Glu^ neurons and heroin seeking behavior, both of which were successfully rescued by chemogenetic activation of RSG^Glu^ neurons (Figures 4U-4W). These results together support that Cck/Cckbr signaling plays a critical role in modulation of RSG^Glu-Cckbr^ neuronal activity and is required for the encoding and retrieval of heroin-associated memories.

### ZI^GABA^, but not ZI^DA^-projecting layer 5 RSG^Glu-Cckbr^ neurons play a key role in context-induced relapse to heroin seeking

Given that the RSC stands as a major hub for interconnectivity among multiple cortical and subcortical brain regions^21^, we sought to identify the circuit-based mechanisms by which RSG^Glu-Cckbr^ neurons control relapse to heroin seeking. To this end, we first detected the effect of silence of the synaptic output of RSG^Glu^ neurons on CIR behavior by expressing the tetanus toxin light chain (TeNT) in these neurons. Suppression of RSG neuronal output significantly decreased CIR behavior (Figures S11A-S11C), supporting circuit mechanisms underlying RSG regulation of CIR.

To determine the efferent targets of RSG^Glu-Cckbr^ neurons, we injected AAV-Cckbr-Cre and AAV-CaMKIIα-DIO-mCherry into RSG (Figures S12A and S12B), which resulted in robust labeling of several cortical and subcortical regions, including anterior cingulate cortex (ACC), anterodorsal nucleus of thalamus (AD), subiculum (Sub) and zona incerta (ZI) (Figure S12C), indicating that these brain regions were innervated by RSG^Glu-Cckbr^ neurons. RSG possesses the typically laminar structure that can be divided into layer 1, layer 2/3, layer 5 and layer 6^37,38^ (see Allen Brain Atlas in Figure S13A). To further identify the circuit composed of laminar projections from RSG to downstream brain regions involved in regulation of CIR, we first mapped the laminar projection profile of RSG^Glu^ neurons by retrograde tracing. We injected scAAV2/R-hSyn-EGFP into ZI, Sub and scAAV2/R-hSyn-mCherry into ACC and AD, respectively (Figures S12D and S12E). We found that neurons in RSG labeled by EGFP or mCherry from Sub or ACC were distributed in both layer 5 and 6, whereas neurons in RSG labeled by EGFP from ZI were exclusivel<u>y</u> distributed in layer 5 (Figures S12D and S12E). RSG layer 5 neurons display more greatly labeling from ZI than that from ACC and Sub (Figure S12G), suggesting that RSG layer 5 neurons preferentially project to ZI. In addition, we also observed retrograde-labeled neurons in RSA. However, the proportion of labeled neurons in RSA from ZI was relatively lower than those from ACC and Sub (Figure S12H), indicating that ZI received less projections from RSA, whereas ACC and Sub received more projections from RSA. Due to the low retrograde tracing efficiency of AAV2/R in AD, we instead injected fluorogold, a strong retrograde tracer, into AD, and observed intense neuronal fluorescence in RSG layer 6 (Figure S12F). Together, these results demonstrate that RSG exhibits layer-distinct projection profile and ZI is one of the primary downstream projecting targets of the RSG layer 5 neurons.

To further determine RSG^Glu-Cckbr^ neurons from which layers project to ZI, we first identified the distributions of RSG^Glu-Cckbr^ neurons in discrete layers by utilizing the Stereo-seq-based spatial transcriptome analysis after CIR test (Figures 5A and 5B). Using dimension reduction and clustering analysis, we defined seven neuronal clusters by the high expression of *Snap25*, including clusters 4, 6, 7, 10, 11, 13 and 15 (Figures 5C, S13B and S13C). The clusters 4, 6, 7, 10 and 11, which highly express *Slc17a7* gene, are considered as glutamatergic neurons and the clusters 13 and 15, which highly express *Gad1* and *Gad2*, are considered as GABAergic neurons (Figure S13D). Interestingly, the spatial distribution of the glutamatergic clusters displayed laminar characteristics, with the clusters 4 and 6 in layer 2/3, the cluster 11 and a minority of cluster 7 in layer 5, and a majority of cluster 7 in layer 6. However, the two GABAergic clusters were scattered across the whole RSG region without layer specificity (Figure 5D). We then analyzed the average expression of *Cckbr* in each glutamatergic cluster and found that cluster 11 (layer 5) expressed the highest level of *Cckbr*, followed by cluster 6 (layer 2/3) and 7 (layer 5 and 6) (Figure 5E). Highly expressing of Cckbr in layer 5 was further confirmed by immunostaining (Figures 5F-5H). We also found that heroin SA training resulted in significantly increased Cckbr protein expression in RSG layer 5 rather than in other layers (Figures 5I and S14A-S14C). Using the marker genes for distinct cortical layers^37^, we can accurately recognize the layer 2/3, layer 5 and layer 6 of RSG (Figures 5J and S13E). Clustering and marker gene analysis revealed the transcriptional diversity across the different layers (Figures S13F-S13H). Next, we analyzed the expression of several IEGs chosen from our snRNA-seq data in different RSG layers. It was found that while *Homer1*, *Egr1* and *Arhgef3* were expressed highly in layers 5 and 6, *Dusp1* and *Arc* were higher expressed only in layer 5 (Figure 5K). Among these IEGs, *Homer1*, *Arhgef3*, *Arc* and *Dusp1* have been shown to be involved in synaptic plasticity, learning and memory as well as drug addiction^39–46^. These results together suggested that layer 5 RSG^Glu-Cckbr^ neurons may play an important role in the modulation of CIR behavior through projections to ZI.

**Figure 5.**
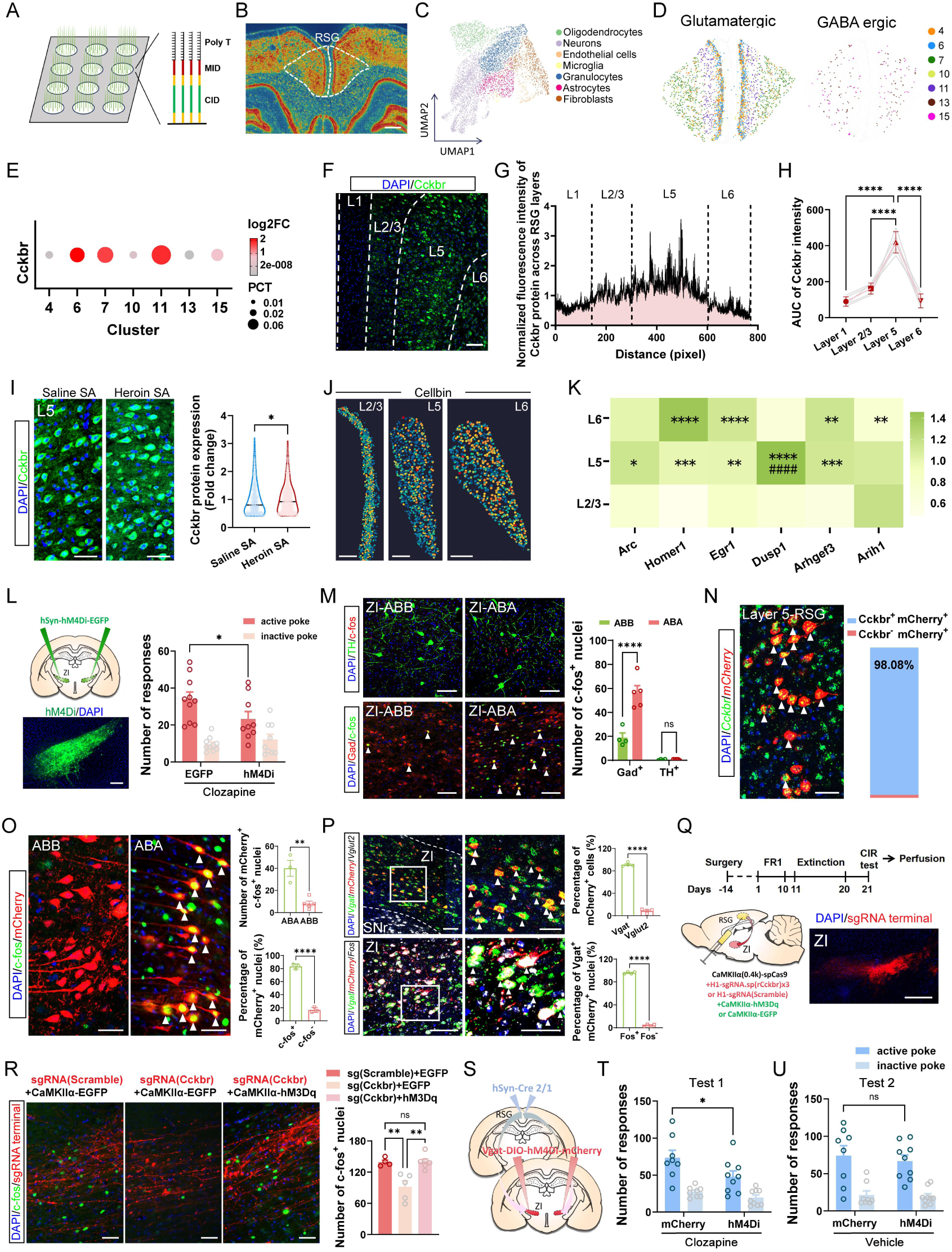
Layer 5 RSG^Glu-Cckbr^-ZI^GABA^ circuit plays a key role in retrieval of contextual heroin memory. **(A)** Schematic showing the sequencing chip of stereo-seq technology. **(B)** Visualization of the spatial transcriptome of the coronal brain slice containing RSG region. Scale bars, 300 μm. **(C)** Clustering analysis of RSG cells visualized by Uniform manifold approximation and projection (UMAP) dimensional reduction. **(D)** Spatial distribution of different clusters of glutamatergic and GABAergic neurons in RSG. **(E)** Dotplot showing the *Cckbr* mRNA expression in different clusters of RSG glutamatergic and GABAergic neurons. **(F)** Representative image showing the expression of Cckbr protein in RSG. Scale bars, 100 μm. **(G)** Normalized fluorescence intensity of Cckbr protein across the different layers of RSG. **(H)** Area under curve of the fluorescence intensity of Cckbr protein in different layers of RSG (*n* = 5). One-way ANOVA (F_(3, 16)_ = 72.32, *p* < 0.0001) followed by Tukey’s post hoc test, \*\*\*\**p* < 0.0001. **(I)** Left: representative images showing the expression of Cckbr protein in RSG layer 5 of rats in Saline SA and Heroin SA groups. Scale bars, 50 μm. Right: average expression level of Cckbr protein in RSG layer 5 of Saline SA (*n* = 3) *vs* Heroin SA (*n* = 3) rats. Mann-Whitney test, \**p* < 0.05. **(J)** Recognition and separation of different layers in RSG. Scale bars, 200 μm. **(K)** Heatmap showing the differential IEGs expression in RSG layer 2/3, layer 5 and layer 6. Multiple Mann-Whitney test followed by False Discovery Rate (FDR) post test, \**p* < 0.05, \*\**p* < 0.01, \*\*\**p* < 0.001, \*\*\*\**p* < 0.0001 *vs* L2/3, ^####^*p* < 0.0001 *vs* L6. **(L)** Left: schematic of the viral strategy for chemogenetic inhibition of ZI neurons and the representative image showing the hM4Di expression in ZI. Scale bars, 100 μm. Right: number of responses of rats in EGFP control group (*n* = 10) *vs* hM4Di group (*n* = 9). Two-way ANOVA (F_(1,34)_ = 1.635, *p* = 0.2096) followed by Sidak’s post hoc test, \**p* < 0.05. **(M)** Left: representative images showing the expression of TH and c-fos (top) or Gad and c-fos (bottom) in ZI of rats in ABB group and ABA group. Scale bars, 100 μm. Right: number of c-fos-positive cells in TH^+^ or Gad^+^ neurons in ZI of ABB group (*n* = 4) *vs* ABA group (*n* = 5). Two-way ANOVA (F_(1,14)_ = 25.68, *p* < 0.001) followed by Sidak’s post hoc test, \*\*\*\**p* < 0.0001, ns, no significant difference. **(N)** Left: representative image showing the co-localization of *Cckbr* and the mCherry-labeled ZI-projecting neurons in RSG layer 5. Scale bars, 50 μm. Right: percentage of Cckbr^+^ and Cckbr^-^ cells in mCherry^+^ neurons in RSG layer 5 (*n* = 3). **(O)** Left: representative images showing the expression of mCherry and c-fos in RSG layer 5 of rats in ABB group and ABA group. Scale bars, 50 μm. Right: number of mCherry^+^ c-fos^+^ neurons in RSG layer 5 of ABB group (*n* = 5) *vs* ABA group (*n* = 3) and percentage of Fos^+^ and Fos^-^ nuclei in mCherry^+^ cells in RSG layer 5 in ABA group. Unpaired t test, \*\**p* < 0.01, \*\*\*\**p* < 0.0001. **(P)** Top: representative images showing the co-localization of *Vgat*, *Vglut2* and *mCherry* in ZI, and percentage of mCherry-positive cells in Vglut2^+^ and Vgat^+^ neurons in ZI (*n* = 4). Unpaired t test, \*\*\*\**p* < 0.0001. Bottom: representative images showing the co-localization of *Vgat*, *mCherry* and *Fos* in ZI after context-induced relapse, and percentage of Fos^+^ and Fos^-^ nuclei in Vgat^+^ mCherry^+^ cells in ZI after context-induced relaspe (*n* = 4). Scale bars, 100 μm and 50 μm. Unpaired t test, \*\*\*\**p* < 0.0001. **(Q)** Schematic showing the training and perfusion schedule, the viral strategy for Cckbr knockout and chemogenetic activation of RSG glutamatergic neurons and the representative image of RSG axon terminals in ZI. Scale bars, 500 μm. **(R)** Left: representative images showing the c-fos expression in ZI adjacent to the axon terminals of RSG glutamatergic neurons in rats of control, Cckbr knockdown and Cckbr knockdown with hM3Dq groups after context-induced relapse. Scale bars, 50 μm. Right: number of c-fos-positive neurons in ZI of rats in control (*n* = 4), Cckbr knockdown (*n* = 5) and Cckbr knockdown with hM3Dq (*n* = 6) groups after context-induced relapse. One-way ANOVA (F(_2, 12)_ = 12.30, *p* < 0.01) followed by Tukey’s post hoc test, \*\**p* < 0.01, ns, no significant difference. **(S)** Schematic of the viral strategy for chemogenetic inhibition of RSG^Glu-Cckbr^-ZI^GABA^ circuit. **(T)** Number of responses in mCherry control group (*n* = 8) and hM4Di group (*n* = 9) during test 1 with clozapine injection (i.p.). Two-way ANOVA (F_(1,30)_ = 6.145, *p* < 0.05) followed by Sidak’s post hoc test, \**p* < 0.05. **(U)** Number of responses in mCherry control group (*n* = 8) and hM4Di group (*n* = 9) during test 2 with vehicle injection. Two-way RM ANOVA (F_(1,30)_ = 0.3091, *p* = 0.5824) followed by Sidak’s post hoc test. ns, no significant difference.

ZI, a large subthalamic region, is regarded as a potential integrative node for global behavioral modulation^47^ and plays a critical role in control of motivated behaviors^48–50^. To investigate the role of ZI in CIR, we examined whether ZI was activated by CIR by using c-fos immunostaining. We found that CIR robustly activated ZI neurons (Figure S15A). Chemogenetic inhibition of ZI neurons attenuated context-induced relapse to heroin (Figure 5L). ZI is a heterogeneous nucleus that expresses major GABAergic neurons and it also expresses dopaminergic neurons^51,52^. Using Tyrosine hydroxylase (TH) and Gad as markers for dopaminergic and GABAergic neurons, we found that the CIR activated GABAergic rather than dopaminergic neurons (Figure 5M), suggesting that ZI GABAergic (ZI^GABA^) neurons are pivotally implicated in CIR behaviors.

Next, we asked whether ZI^GABA^ neuron-projecting layer 5 RSG^Glu-Cckbr^ neurons (RSG^Glu-Cckbr^-ZI^GABA^ circuit) played a key role in the retrieval of heroin-associated memories, leading to relapse to heroin seeking. We first determined whether the ZI-projecting layer 5 RSG^Vglut1^ neurons were Cckbr positive glutamatergic neurons by use of a combination of retrograde tracing and FISH. We confirmed that these neurons were Cckbr-positive glutamatergic neurons (Figures 5N, S15B and S15C). We also found that these neurons were significantly activated by CIR (Figure 5O). Using trans-synaptic anterograde tracing combined with FISH or TH immunostaining, we found that RSG neurons form direct synaptic connections with ZI GABAergic, but not dopaminergic or glutamatergic neurons (Figures 5P and S15D-S15F), and that CIR induced robust activation of the RSG-innervated ZI GABAergic neurons (Figure 5P).

Next, we detected the role of Cckbr in modulating activity of RSG^Glu-Cckbr^-ZI^GABA^ circuit during CIR by knockdown of Cckbr in RSG^Glu-Cckbr^ neurons with CRISPR-Cas9 (Figure 5Q). We found that CIR-induced activation of ZI^GABA^ neurons was significantly impaired by knockdown of Cckbr in RSG^Glu-Cckbr^ neurons, compared to the rats in the control group. However, such effect evoked by Cckbr knockdown could be rescued by chemogenetic activation of RSG^Glu^ neurons (Figure 5R), indicating the importance of Cckbr for regulation of the RSG^Glu-Cckbr^-ZI^GABA^ circuit. Accordingly, chemogenetic inhibition of RSG^Glu-Cckbr^-ZI^GABA^ circuit also significantly decreased CIR behavior (Figures 5S-5U). Together, these results support that the RSG^Glu-Cckbr^-ZI^GABA^ circuit plays a key role in regulation of CIR behavior.

### Anterodorsal nucleus of thalamus controls the activity of ZI^GABA^-projecting layer 5 RSG^Glu-Cckbr^ neurons by release of Cck

We next sought to identify the upstream Cckergic projections that release of Cck for activating RSG^Glu-Cckbr^-ZI^GABA^ circuit. To this end, we first conducted rabies virus (RV)-mediated monosynaptic retrograde tracing. Two AAVs expressing Cre-dependent avian-specific retroviral receptor (TVA) fused with mCherry and rabies virus glycoprotein (RVG) were injected into RSG, and the AAV2/R-EF1α-Cre was injected into ZI. Two weeks later, G-deleted EnvA-pseudotyped RV (RV-EnvA-ΔG-EGFP) was delivered into RSG (Figure 6A). The starter neurons that co-expressed mCherry and EGFP could be detected in the RSG layer 5 (Figure 6B). Next, whole brain mapping of the upstream projections was conducted using brain clearing and 3D imaging (Figure 6A). We found that the ZI-projecting layer 5 RSG^Glu-Cckbr^ neurons received internally ipsilateral and contralateral projections from RSG and external innervation from several cortical and subcortical brain regions, including RSA, ACC, Secondary Motor Cortex (M2), AD, Sub, Claustrum (Cla) and Horizontal Diagonal Band (HDB) (Figures 6B and 6C). Among these upstream regions, ipsilateral RSG showed stronger internal projections than contralateral RSG (Figure 6D), and ACC and RSA contributed more external innervation compared to other regions (Figure 6E).

**Figure 6.**
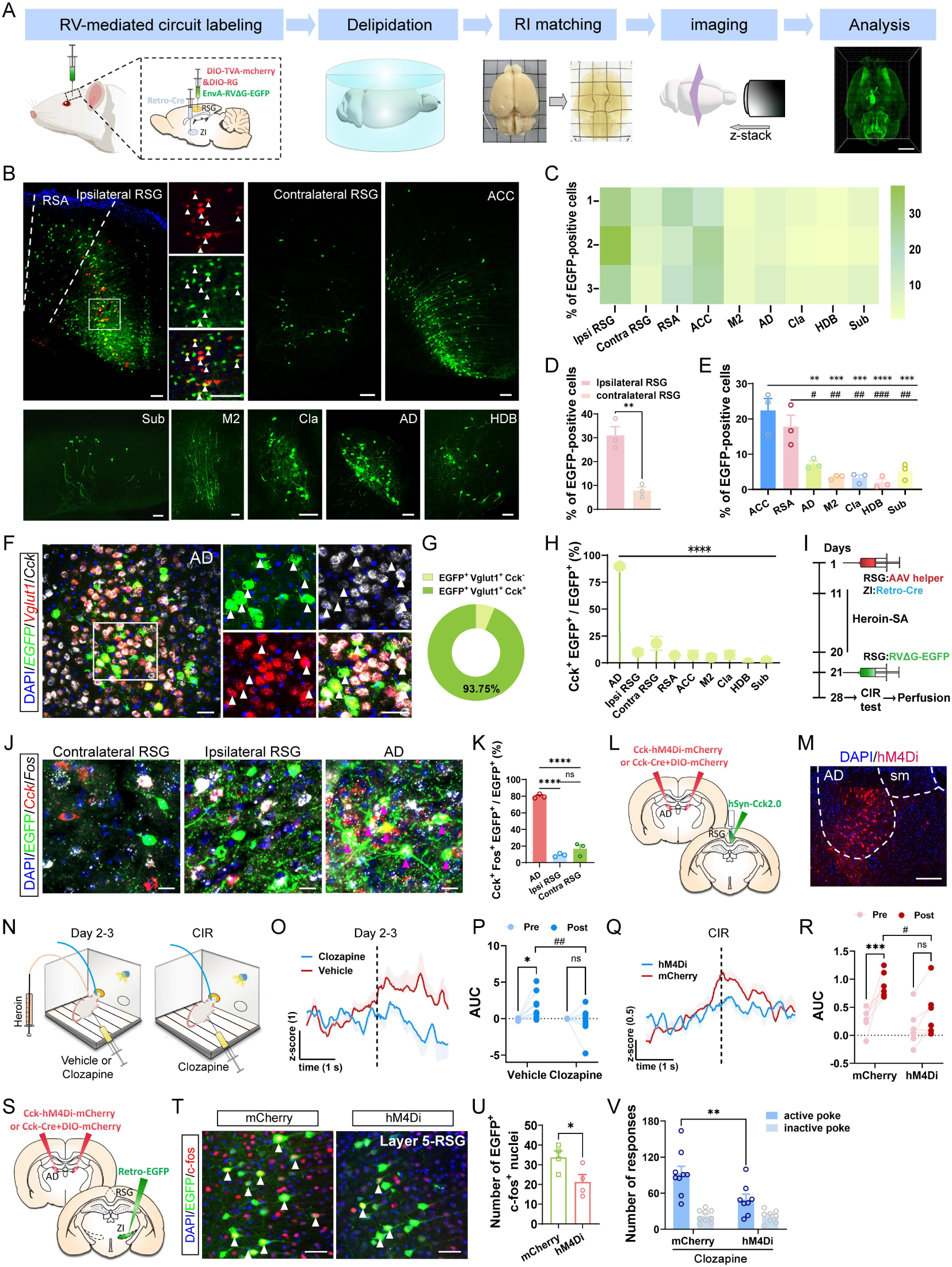
Anterodorsal nucleus of thalamus controls the activity of ZI-projecting layer 5 RSG^Glu-Cckbr^ neurons by release of Cck. **(A)** Schematic showing the viral strategy and brain tissue clearing for the whole-brain mapping of the upstream innervation to ZI-projecting layer 5-RSG^Glu-Cckbr^ neurons. Scale bars, 5000 μm. **(B)** Representative images showing the starter cells in RSG and the retrograde-labeled neurons in upstream brain regions. Scale bars, 100 μm. ACC: anterior cingulate cortex, AD: anterodorsal nucleus, Sub: subiculum, M2: secondary motor cortex, Cla: claustrum, HDB: horizontal diagonal band. **(C)** Heatmap of the percentage of EGFP-labeled neurons in every upstream brain region in individual rat (*n* = 3, % of total EGFP-positive neurons). **(D)** Percentage of the EGFP-labeled neurons in internal afferent from ipsilateral *vs* contralateral RSG (*n* = 3). Unpaired t test, \*\**p* < 0.01. **(E)** Percentage of the EGFP-labeled neurons in external afferent brain regions (*n* = 3). One-way ANOVA (F_(6, 14)_ = 16.72, *p* < 0.0001) followed by Tukey’s post hoc test, \*\**p* < 0.01, \*\*\**p* < 0.001, \*\*\*\**p* < 0.0001 vs ACC; ^#^*p* < 0.05, ^##^*p* < 0.01, ^###^*p* < 0.001 vs RSA. **(F)** Representative images showing the co-localization of *EGFP*, *Vglut1* and *Cck* in AD. Scale bars, 50 μm. **(G)** Percentage of Cck-positive and negative cells in EGFP^+^ Vglut1^+^ neurons in AD (*n* = 3). **(H)** Percentage of EGFP-labeled Cck-positive neurons in AD and other afferent brain regions (*n* = 3). One-way ANOVA (F_(8,18)_ = 72.13, *p* < 0.0001) followed by Tukey’s post hoc test, \*\*\*\**p* < 0.0001 vs AD. **(I)** Schematic showing the schedule for virus injection, heroin SA training, context-induced relapse and perfusion. **(J)** Representative images showing the co-localization of EGFP, *Fos* and *Cck* in contralateral RSG, ipsilateral RSG, and AD. Scale bars, 20 μm. **(K)** Percentage of Cck^+^ Fos^+^ EGFP^+^ neurons in EGFP-labeled cells in AD, contralateral and ipsilateral RSG. One-way ANOVA (F_(2,6)_ = 189.5, *p* < 0.0001) followed by Tukey’s post hoc test, \*\*\*\**p* < 0.0001. **(L)** Schematic showing the viral strategy for chemogenetic inhibition of AD Cckergic neurons and the fiber photometry recording of Cck in RSG. **(M)** Representative image of the expression of hM4Di in AD. Scale bars, 200 μm. **(N)** Schematic showing the schedule for fiber photometry recording. **(O)** Average Cck signals in RSG during early (day 2-3) phase of heroin SA training in hM4Di group with clozapine or vehicle injection. Vertical dashed line indicates the onset of active poke behaviors. **(P)** Area under curve of the Cck signals in clozapine (*n* = 8) and vehicle (*n* = 10) groups during early (day 2-3) phase of heroin SA training. Two-way RM ANOVA (F_(1,16)_ = 6.174, *p* < 0.05) followed by Sidak’s post hoc test, \**p* < 0.05. ^##^*p* < 0.01. **(Q)** Average Cck signals during context-induced relapse in hM4Di and mCherry groups. Vertical dashed line indicates the onset of active poke behaviors. **(R)** Area under curve of the Cck signals in hM4Di (*n* = 7) and mCherry (*n* = 6) group during context-induced relapse. Two-way RM ANOVA (F_(1,11)_ = 4.824, *p* = 0.0504) followed by Sidak’s post hoc test, \*\*\**p* < 0.001, ^##^*p* < 0.05. **(S)** Schematic of the viral strategy for chemogenetic inhibition of AD Cckergic neurons and labeling of ZI-projecting neurons in RSG layer 5. **(T)** Representative images showing the expression of EGFP and c-fos in RSG layer 5 of rats in mCherry control group and hM4Di group during context-induced relapse. Scale bars, 50 μm. **(U)** Number of EGFP^+^ c-fos^+^ neurons in RSG layer 5 of mCherry group (*n* = 4) vs hM4Di group (*n* = 4) during context-induced relaspe. Unpaired t test, \**p* < 0.05. **(V)** Number of responses in mCherry control group (*n* = 9) and hM4Di group (*n* = 8) during context-induced relaspe. Two-way ANOVA (F_(1,30)_ = 7.564, *p* < 0.01) followed by Sidak’s post hoc test, \*\**p* < 0.01.

Next, we explored the Cckergic projections from upstream brain regions by employing FISH combined with RV-mediated retrograde tracing. Interestingly, quantitative analysis revealed that while the ZI-projecting layer 5 RSG^Glu-Cckbr^ neurons received a small amount of internal Cck input from ipsilateral and contralateral RSG and external Cck input from RSA, ACC, M2, Sub and Cla, these neurons received the densest external Cck input from AD (Figures 6F-6H and S16A-S16C). We further examined the CIR-induced activation of the Cckergic projections in the relatively higher three Cck input brain regions (Figure 6I) and found that the proportion of *Fos* expression in *Cck* positive input neurons in AD was dramatically higher than that in ipsilateral and contralateral RSG (Figures 6J and 6K), indicating that CIR strongly activated Cck projections from AD to ZI-projecting layer 5 RSG^Glu-Cckbr^ neurons. Next, we determined the role of RSG-projecting AD Cckergic (AD^Cck^) neurons in heroin SA training and CIR-induced release of Cck, activation of RSG^Glu-Cckbr^ neurons and heroin seeking behavior. We found that chemogenetic inhibition of AD^Cck^ to RSG projections suppressed heroin SA training and CIR-induced augmentation of Cck release (Figures 6L-6R), activation of ZI^GABA^-projecting RSG^Glu-Cckbr^ neurons (Figures 6S-6U) and induction of context-induced relapse to heroin (Figure 6V). Together, these results suggest that AD orchestrates RSG^Glu-Cckbr^-ZI^GABA^ circuit function through release of Cck.

## DISCUSSION

In this study, we define the role of a new top-down pathway from the RSG subregion of the RSC to the ZI in the control of retrieval of long-term opioid-associated memories, leading to relapse behavior. This cortico-subthalamic pathway, which is not as a canonical pathway responsible for encoding drug addictive behaviors as the well-studied reciprocal pathway in the mesocorticolimbic regions, is activated in response to heroin SA training and context-induced relapse to heroin. RSG^Glu-Cckbr^ neurons of the layer 5 exert their influence on relapse behavior through excitation of ZI GABAergic, but not dopaminergic neurons. Cck derived from AD thalamus orchestrates these ZI-projecting RSG^Glu-Cckbr^ neurons. These results elucidate a non-canonical top-down pathway regulating retrieval of long-term opioid-associated memories, leading to relapse behavior.

The high relapse rates in opioid abstinent individuals may be caused by abnormally long-lasting opioid-associated memories^53–55^. Repetitive exposures to abused drugs and withdrawal of drug use can lead to synaptic plasticity, a key cellular mechanism underlying learning and memory. Development of drug addictive behaviors has been proposed to be attributable to an aberrant form of learning and memory^56–59^. Drugs of abuse-induced alteration of dendritic spine structure and rewiring of neuronal circuitry may underlie drug-induced long-term drug-associated memories^60,61^. However, which brain regions are responsible for the encoding, consolidation and retrieval of long-term opioid-associated memories and how these brain regions achieve their goals are still poorly understood.

Several lines of evidence suggest that neocortical areas such as mPFC, ACC, and RSC may be important brain regions that engage in the storage and retrieval of remote memories^62,63^. The RSC is of particularly interesting because of its anatomic connectivity with hippocampus, visuo-sensory and motor cortex and its role in spatial, contextual, and episodic memories^64,65^. RSC is thought to be a critical site for long-term memory storage and retrieval^25,65–68^. Patients with RSC damage or lesion of RSC exhibited severe retrograde amnesia^67,69^. Functional magnetic resonance imaging studies in humans show increased RSC activity during the retrieval of remote spatial^70^, autobiographical memories^71,72^ in healthy subjects and traumatic memories in patients with post-traumatic stress disorder^73^. Our study further reveals that RSG is required for the encoding and retrieval of opioid-associated memories, as ablation or chemogenetic inhibition of RSG^Glu^ neurons abolishes heroin self-administration behavior and context-induced relapse to heroin. Similarly, a previous study also reported that retrieval of morphine conditioned place preference (CPP) memory is accompanied by increasing c-fos expression in RSC^74^. Using snRNA-seq technology, we found that RSG^Vglut1^ neurons express multiple gene transcripts that encode several receptors critically involved in emotion, recognition, learning and memory including cholecystokinin B receptor (*Cckbr*), serotonin receptor 2c and 4 (*Htr2c, Htr4*), glutamate ionotropic receptor AMPA type subunit 1 (*Gria1*), cholinergic receptor muscarinic 3 and nicotinic alpha 5 (*Chrm3, Chrna5*). Employing spatial transcriptomics (Stereo-seq) and immunostaining analysis, we further found that the spatial distribution of the glutamatergic clusters displayed laminar characteristics, with the clusters 4 and 6 in layer 2/3, the cluster 11 and a minority of cluster 7 in layer 5, and a majority of cluster 7 in layer 6, and that RSG^Vglut1^ neurons of layer 5 expressed the highest level of *Cckbr*. RSG Cckbr positive glutamatergic neurons play a pivotal role in encoding and retrieval of opioid-associated memories, since heroin SA training and context-induced relapse resulted in activation of these neurons by upregulation of RSG Cck concentrations, an endogenous ligand of Cckbr. Heroin SA training and context-induced relapse failed to upregulate endogenous ligands for other receptors detected in RSG.

Cck is one of the most abundant cortical neuropeptide^75^ and plays a significant role in synaptic transmission and plasticity within the CNS^76–78^. It has been shown that Cck-positive GABAergic^79–82^ and glutamatergic neurons^34,83–85^ both exist in the neocortex, which enables cortical synaptic plasticity and associative memory formation. Several studies show that cortical Cck-positive glutamatergic and GABAergic neurons participate in the formation of long-term heterosynaptic plasticity^86^ and long-term memory^80,82^. Using gene-encoded neurotransmitter sensors, we demonstrate that heroin SA training and context-induced relapse to heroin evoke a significant increase in Cck rather than other neurotransmitter levels in RSG. Cck plays a key role in encoding of long-term opioid-associated memories via activation of Cckbr in excitatory RSG^Glu^ neurons. Knockdown of Cckbr with shRNA or CRISPR-Cas9 impaired the activation of RSG glutamatergic neurons and robustly decreased heroin SA behavior and context-induced relapse to heroin.

We further reveal that Cck is derived from AD thalamus. The thalamus has been shown to contribute to the persistence of frontal cortical activity during motor preparation, associative learning and working memory maintenance^87–91^. The anterior thalamic nuclei (ATN) have reciprocal connectivity with frontal cortical areas such as RSC and ACC and are necessary for contextual memory encoding and remote memory retrieval^38,92–94^. AD thalamus, a part of the understudied ATN complex, is specifically important for contextual encoding processes, as it directly receives visual input^95^. Using FISH combined with RV-mediated retrograde tracing, we find that ZI-projecting Cck-positive RSG^Glu^ neurons in layer 5 receive predominant Cck projections from the AD thalamus. Chemogenetic inhibition of AD^Cck^ neuron<u>s</u> suppresses heroin SA training and CIR-induced Cck release, activation of ZI^GABA^-projecting RSG^Glu-Cckbr^ neurons and context-induced relapse to heroin. Consistent with a recent study showing that the AD thalamus-RSG circuit is necessary for contextual fear memory encoding^93^, the present study indicates that the AD thalamus-RSG circuit may play an important role in the encoding and retrieval of longer-term opioid-associated memories.

The ZI is a large subthalamic region with extensive connections throughout the brain^96^, and is regarded as a potential integrative node for global behavioral modulation^47^. Substantial evidence supports that ZI plays a critical role in encoding appetitive motivation and controlling motivated behaviors^48–50^. ZI is also reported to be required for remote recall of conditioned fear memory^97^. ZI is a largely inhibitory nucleus comprised of heterogeneous groups of cells. It expresses multiple molecular markers such as parvalbumin (PV), tyrosine hydroxylase and glutamate^51,52^. Activation of ZI GABAergic neurons exhibits strong repetitive and progressive self-stimulation and predatory hunting^48,49,98^, whereas activation of ZI dopaminergic neurons encodes motivational vigor in food seeking^50^. In this study, by using a combination of the snRNA-seq, Stereo-seq analysis and viral tracing techniques, we uncover that RSG glutamatergic neurons exhibit laminar diversity in expression of Cckbr and projection profiles. In RSG, layer 5 glutamatergic neurons express higher Cckbr relative to those in other layers and these RSG Cckbr-positive neurons mainly project to ZI GABAergic, but not dopaminergic neurons. Exposure of rats to the heroin self-administration context after extinction robustly activates ZI GABAergic, but not dopaminergic neurons. Chemogenetic inhibition of RSG-innervated ZI^GABA^ neurons could disrupt context-induced relapse to heroin. Thus, we proposed that RSG^Glu-Cckbr^-ZI^GABA^ circuit is crucially involved in the transformation of opioid-associated memories to heroin seeking motivation.

In sum, the present study provides insight into a mesocorticolimbic pathway-independent circuit mechanism underlying the encoding and retrieval of opioid-associated memories and may help guide the development of new therapeutic strategies to prevent relapse to opioid seeking.

## ACKNOWLEDGMENT

This research was supported by the Major Project of the Science and Technology Innovation 2030 of China (STI2030-Major Projects 2021ZD0202900 to J.-G.L., 2021ZD0203500 to Y.-J.W.), the National Natural Science Foundation of China (82030112 to J.-G.L., 82571743 to Y.-J.W., 82301677 to R.-S.C.), the Science and Technology Commission of Shanghai Municipality (23ZR1474900 to Y.-J.W.), the Key R&D Program and the Taishan Scholars Program of Shandong Province, China (2024CXPT029 to Y.-J.W.).

## AUTHOR CONTRIBUTION

Z.-Q.W., Y.-J.W., and J.-G.L. designed the experiments. J.L., and R.-S.C. performed the experiments and statistical analysis with the help of R.-F.Y., G.-Y.Z., Z.-H.L., S.W., Y.-Q.S., X.-Q.Z., and C.X. The manuscript was written by R.-S.C., and J.L., and was revised by Y.-J.W., and J.-G.L.

## STAR METHODS

KEY RESOURCES TABLE.

KEY RESOURCES TABLE
| REAGENT or RESOURCE | SOURCE | IDENTIFIER |
| --- | --- | --- |
| Antibodies |  |  |
| Rabbit-anti-c-fos | Synaptic systems | Cat# 226008; RRID: AB_2891278 |
| Mouse-anti-c-fos | Abcam | Cat# ab208942; RRID: AB_2747772 |
| Rabbit-anti-TH | Millipore | Cat# ab152; RRID: AB_390204 |
| Mouse-anti-CaMKIIα | Abcam | Cat# ab22609; RRID: AB_447192 |
| Mouse-anti-Gad | Abcam | Cat# ab26116; RRID: AB_448990 |
| Rabbit-anti-Cckbr | Alomone labs | Cat# ACR-042; RRID: AB_2039803 |
| Goat-anti-rabbit (Alexa Fluor 488) | ThermoFisher Scientific | Cat# A-11008; RRID: AB_143165 |
| Goat-anti-rabbit (Alexa Fluor 647) | ThermoFisher Scientific | Cat# A-21244; RRID: AB_2535812 |
| Goat-anti-mouse (Alexa Fluor 555) | ThermoFisher Scientific | Cat# A-21422; RRID: AB_141822 |
| Bacterial and virus strains |  |  |
| AAV2/9-CaMKIIα-mCherry | BrainVTA | Cat# PT-0108 |
| AAV2/9-CaMKIIα-GCaMP6s | BrainVTA | Cat# PT-0110 |
| AAV2/9-CaMKIIα-DIO-GCaMP6s | BrainVTA | Cat# PT-0090 |
| AAV2/9-CaMKIIα-DIO-mCherry | BrainVTA | Cat# PT-1167 |
| AAV2/9-CaMKIIα-DO-GCaMP6s | BrainVTA | Cat# PT-4451 |
| AAV2/9-CaMKIIα-DO-mCherry | BrainVTA | Cat# PT-4454 |
| AAV2/9-CaMKIIα-hM4Di-mCherry | BrainVTA | Cat# PT-0017 |
| AAV2/9-CaMKIIα-hM3Dq-mCherry | BrainVTA | Cat# PT-0016 |
| AAV2/9-CaMKIIα-hM3Dq-EGFP | BrainVTA | Cat# PT-0525 |
| AAV2/9-CaMKIIα-EGFP | BrainVTA | Cat# PT-0290 |
| AAV2/9-CaMKIIα-DIO-hM4Di-mCherry | BrainVTA | Cat# PT-1143 |
| AAV2/9-CaMKIIα-Cre | BrainVTA | Cat# PT-0220 |
| AAV2/9-Vgat-mCherry | BrainVTA | Cat# PT-0325 |
| AAV2/9-Vgat-GCaMP6s | BrainVTA | Cat# PT-5608 |
| AAV2/9-Vgat-DIO-GCaMP6s | BrainVTA | Cat# PT-6173 |
| AAV2/9-Vgat-DIO-mCherry | BrainVTA | Cat# PT-3225 |
| AAV2/9-Vgat-DIO-hM4Di-mCherry | BrainVTA | Cat# PT-0618 |
| AAV2/9-EF1α-DIO-TeNT-EGFP | BrainVTA | Cat# PT-2434 |
| AAV2/9-EF1α-DIO-taCasp3-TEVp-EGFP | BrainVTA | Cat# PT-1230 |
| AAV2/9-EF1α-DIO-EGFP | BrainVTA | Cat# PT-0795 |
| AAV2/9-EF1α-DIO-mCherry | BrainVTA | Cat# PT-0013 |
| AAV2/9-hSyn-iGluSnFR | BrainVTA | Cat# PT-1140 |
| AAV2/9-hSyn-5-HT3.5 | BrainVTA | Cat# PT-2366 |
| AAV2/9-hSyn-Cck2.0 | BrainVTA | Cat# PT-1354 |
| AAV2/9-hSyn-Ach3.0 | BrainVTA | Cat# PT-1335 |
| AAV2/9-hSyn-EGFP | BrainVTA | Cat# PT-1990 |
| AAV2/9-hSyn-hM4Di-EGFP | BrainVTA | Cat# PT-0153 |
| AAV2/9-Cckbr-Cre-EGFP | BrainVTA | Cat# PT-5868 |
| AAV2/9-Cck-hM4Di-mCherry | BrainVTA | Cat# PT-13162 |
| AAV2/9-Cck-Cre | BrainVTA | Cat# PT-6012 |
| AAV2/1-hSyn-Cre | BrainVTA | Cat# PT-0136 |
| AAV2/R-EF1α-Cre | BrainVTA | Cat# PT-0888 |
| scAAV2/R-hSyn-mCherry | BrainVTA | Cat# PT-3975 |
| scAAV2/R-hSyn-EGFP | BrainVTA | Cat# PT-2315 |
| AAV2/9-CaMKIIα-shRNA(Cckbr)-mCherry | OBiO | Cat# Y34571 |
| AAV2/9-CaMKIIα-shRNA(Scramble)-mCherry | OBiO | Cat# Y13971 |
| AAV2/9-CaMKIIα(0.4k)-spCas9 | Shanghai Taitool Bioscience | Cat# S1643-9 |
| AAV2/9-H1-sgRNA(Scramble)-hSyn-mCherry | Shanghai Taitool Bioscience | N/A |
| AAV2/9-H1-sgRNA(rCckbr)×3-hSyn-mCherry | Shanghai Taitool Bioscience | N/A |
| CVS-EnvA-RV-ΔG-EGFP | Brain Case | Cat# RV-CVS-EnvA461 |
| AAV2/9-EF1α-DIO-N2cG | Brain Case | Cat# BC0442 |
| AAV2/9-EF1α-DIO-F2A-mCherry-TVA | Brain Case | Cat# BC0061 |
| Chemicals, peptides, and recombinant proteins |  |  |
| Clozapine | Sigma | Cat# C6305 |
| Zoletil | Virbac | N/A |
| L-365260 | MCE | Cat# HY-106840 |
| DAPI | Beyotime | Cat#: C1005 |
| Critical commercial assays |  |  |
| RNAScope Multiplex Fluorescent Reagent Kit V2 Assay | ACD | Cat# 323100 |
| RNAScope Probe-Rn-Slc32a1 | ACD | Cat# 424541 |
| RNAScope Probe-Rn-Slc17a7 | ACD | Cat# 317001 |
| RNAScope Probe-Rn-Slc17a6-C2 | ACD | Cat# 317011-C2 |
| RNAScope Probe-Rn-Cck-C2 | ACD | Cat# 532851-C2 |
| RNAScope Probe-Rn-Cckbr-C2 | ACD | Cat# 312001-C2 |
| RNAScope Probe-Rn-Fos-C3 | ACD | Cat# 403591-C3 |
| RNAScope Probe-mCherry-C3 | ACD | Cat# 431201-C3 |
| RNAScope Probe-EGFP-O4-C3 | ACD | Cat# 538851-C3 |
| Software and algorithms |  |  |
| GraphPad Prism 9.5.1 | GraphPad Software | https://www.graphpad.com |
| Zen system | Zeiss | https://www.zeiss.com/microscopy/us/home.html |
| FV1000 | FV1000 | https://www.olympus-global.com/ |
| ImageJ | National Institute of Health | https://imagej.net/ij/download.html |
| R | R Project for Statistical Computing | RRID: SCR_001905 |

## RESOURCE AVAILABILITY

### Lead contact

Further information and requests for resources and reagents should be directed to and will be fulfilled by the lead contact, Jing-Gen Liu

### Materials availability

This study did not generate new unique reagents.

### Data and code availability

This paper does not report original code. The raw sequence data reported in this paper have been deposited in the Genome Sequence Archive (Genomics, Proteomics & Bioinformatics 2025) in National Genomics Data Center (Nucleic Acids Res 2025), China National Center for Bioinformation / Beijing Institute of Genomics, Chinese Academy of Sciences (GSA: CRA033670) that are publicly accessible at https://ngdc.cncb.ac.cn/gsa as of the date of publication. Data reported in this paper will be shared by the lead contact upon request. Any additional information required to reanalyze the data reported in this paper is available from the lead contact upon request.

## EXPERIMENT MODEL AND SUBJECT DETAILS

### Animals

Male Sprague-Dawley rats weighing 250-280 g were obtained from Shanghai Laboratory Animal Center, Chinese Academy of Sciences, and were housed in ventilated micro-isolator cages under constant conditions (temperature: 23 ± 2 °C, humidity: 50% ± 5% and light/dark cycle: 12 h) with food and water ad libitum. Rats were housed individually after jugular surgery. All husbandry and experiments complied with the National Institutes of Health Guide for the Care and Use of Laboratory Animals and approved by the Institute Animal Care and Use Committee at Shanghai Institute of Materia Medica, Chinese Academy of Sciences.

## METHOD DETAILS

### Stereotaxic surgery

Rats were anesthetized by Zoletil (50 mg/kg, i.p.) and subsequently placed on the stereotaxic apparatus. Standard procedure was used to expose the skull and locate the position of RSG, RSA, ZI, ACC, Sub or AD. Stereotaxic coordinates employed for viral delivery, fiber implantation or cannula implantation were as follows: RSG (AP: –4.0 mm, ML: ±0.40 mm, DV: –2.00 mm), RSA (AP: –4.0 mm, ML: ±1.00 mm, DV: –1.00 mm), ZI (AP: –3.40 mm, ML: ±2.35 mm, DV: –7.50 mm), AD (AP: –1.80 mm, ML: ±1.40 mm, DV: –4.80 mm), ACC (AP: +1.00 mm, ML: ±0.60 mm, DV: –3.00 mm), Sub (AP: –5.20 mm, ML: ±1.60 mm, DV: –3.60 mm). The viral vectors were delivered into the target regions using a micro-infusion pump (Harvard Apparatus, Holliston, MA, USA) at a rate of 300 nl/min and the syringe remained in situ for 7 min to allow the diffusion of the virus.

For fiber photometry recordings, viral vectors of AAV2/9-CaMKIIα-GCaMP6s, AAV2/9-Vgat-GCaMP6s, AAV2/9-hSyn-Cck2.0, AAV2/9-hSyn-Ach3.0, AAV2/9-hSyn-5-HT3.5, AAV2/9-hSyn-iGluSnFR, AAV2/9-Cckbr-Cre combined with AAV2/9-CaMKIIα-DIO-GCaMP6s, AAV2/9-CaMKIIα-shRNA(Cckbr)-mCherry combined with AAV2/9-CaMKIIα-GCaMP6s, AAV2/9-Cckbr-Cre combined with AAV2/9-CaMKIIα-DO-GCaMP6s, AAV2/9-Cckbr-Cre combined with AAV2/9-Vgat-DIO-GCaMP6s were injected unilaterally into the target regions followed by optic fiber implantation. The implanted optic fiber was fixed by dental cement and stainless-steel screws.

For chemogenetic manipulation, viral vectors of AAV2/9-hSyn-hM4Di-EGFP, AAV2/9-CaMKIIα-hM3Dq-mCherry, AAV2/9-CaMKIIα-hM4Di-mCherry, AAV2/9-Cckbr-Cre combined with AAV2/9-CaMKIIα-DIO-hM4Di-mCherry, AAV2/9-Cckbr-Cre combined with AAV2/9-Vgat-DIO-hM4Di-mCherry, AAV2/1-hSyn-Cre, AAV2/9-Vgat-DIO-hM4Di-mCherry and AAV2/9-Cck-hM4Di-mCherry were injected bilaterally into the target regions.

For caspase-induced apoptosis and synaptic silencing, viral vectors of AAV2/9-CaMKIIα-Cre combined with AAV2/9-EF1α-DIO-taCasp3-TEVp-EGFP or AAV2/9-EF1α-DIO-TeNT-EGFP were injected bilaterally into RSG.

For AAV-mediated anterograde tracing, viral vectors of AAV2/9-Cckbr-Cre combined with AAV2/9-CaMKIIα-DIO-mCherry were injected unilaterally into RSG. For AAV-mediated retrograde tracing, viral vectors of AAV2/Retro-hSyn-EGFP and AAV2/Retro-hSyn-mCherry were injected into ZI, ACC, AD and Sub. For fluorogold-mediated retrograde tracing, fluorogold was injected into AD. For trans-synaptic anterograde tracing of RSG-ZI^GABA^ circuit, AAV2/1-hSyn-Cre and AAV2/9-EF1α-DIO-mCherry were injected unilaterally into RSG and ZI, respectively. The histological detection and imaging were conducted at least 4 weeks after surgery.

For RV-mediated retrograde tracing, the viral vectors of AAV2/9-EF1α-DIO-TVA-mCherry combined with AAV2/9-EF1α-DIO-RG and AAV2/Retro-hSyn-Cre were injected unilaterally into RSG and ZI, respectively. 2 weeks later, the vectors of EnvA-RVΔG-EGFP were injected into RSG followed by the histological detection and imaging 1 week later.

For shRNA-mediated mRNA knockdown and CRISPR-Cas9-mediated gene editing, the viral vectors of AAV2/9-CaMKIIα-shRNA(Cckbr)-mCherry or AAV2/9-CaMKIIα(0.4k)-spCas9 combined with H1-sgRNA(rCckbr)×3-hSyn-mCherry and AAV2/9-CaMKIIα-hM3Dq-EGFP were injected bilaterally into RSG. The behavioral training began 2 or 4 weeks after surgery, according to the specific experimental design.

For nucleus microinjection, the cannula was implanted bilaterally 1mm above the RSG region with 14° tilted and was affixed to the skull by dental cement and stainless-steel screws.

### Fiber photometry recording

The fiber photometry recording of signals for Ca^2+^ and neurotransmitter sensors was performed at least 3 weeks after surgery. Before the start of recordings, the pre-implanted fiber was connected to the Multichannel Recording System 810 (RWD Life Science Co. Ltd., Shenzhen, China) or FiberOptoMeter (ThinkerTech Co. Ltd., Nanjing, China) via an optical fiber patch cord (200 μm, 0.37 NA, Inper). The signal was recorded with laser sources of 410 nm and 470 nm at a power of 20-40 μW. Rats were trained for heroin SA under FR1 schedule unless otherwise stated.

For baseline recording, rats were allowed to freely explore the context A for 30 min every day for 3 consecutive days for habituation. The baseline signals of Ca^2+^ or neurotransmitters were recorded on the third day in context A. Then the rats were subjected to heroin SA training.

For recording during early phase of heroin SA, rats were trained for heroin SA on the first day and were introduced into context A and connected to the recording system on day 2 and 3. The tonic signals of freely behaving rats and the phasic signals accompanied by active poke behaviors and heroin infusions were all recorded.

For recording during extinction training, rats were trained for heroin SA for 10 days in context A followed by at least 7-day extinction training in context B. On the last day of extinction training, rats were introduced into context B and the recording system was connected. The tonic signals of freely behaving rats were recorded.

For recording during context-induced relapse test, rats were trained for heroin SA for 10 days in context A followed by at least 7-day extinction training in context B. On the day of relapse test, rats were re-introduced into context A and the tonic signals of freely behaving rats and phasic signals accompanied by active poke behaviors were all recorded.

The data were analyzed and plotted with Multichannel Fiber Photometry Software by RWD or TrippleMulti_6A Software by ThinkerTech. To compare the changes of signals for Ca^2+^ or neurotransmitter sensors accompanied by active nose poke, the area under curve (AUC) of z-score was compared between pre and post of the onset of active poke behavior. To compare the event frequency of the tonic signals, 4 times the mean absolute deviation (4×MAD) of the ΔF/F of baseline signals was used as a threshold for events counting in both signals of baseline and experimental sessions. To compare the amplitude of the tonic signals, the average ΔF/F of signals during baseline and experimental sessions were calculated and analyzed.

### Chemogenetic manipulation

The behavioral training started at least 3 weeks after delivery of viral vectors. Before behavioral training, jugular catheterization surgery was conducted and rats were allowed to recover for 1 week.

For chemogenetic manipulation during context-induced relapse test, rats were trained for heroin SA under fixed ratio (FR) 1 for 10 days in context A followed by at least 7-day extinction training in context B. For the test 1 session, 24 h after the last extinction training session, rats were injected with clozapine i.p. 30 min before the beginning of context-induced relapse test, then rats were re-introduced into context A and the drug seeking behavior was tested for 90 min. Subsequently, rats were maintained in their homecage for 7 days before the test 2 session. During test 2, vehicle was injected i.p. 30 min before re-introducing rats into context A for the 90 min behavioral task.

For chemogenetic activation during extinction training, rats were trained for heroin SA for 10 days in context A followed by at least 7-day extinction training in context B. On the last extinction training session, rats were injected with clozapine i.p. 30 min before being introduced into context B for 90 min behavioral test.

### Caspase-mediated apoptosis

3 weeks after the viral vector injection, jugular catheterization surgery was conducted and rats were allowed to recover for 1 week. Then rats were trained for heroin SA under FR1 (3 days)-FR2 (3 days)-FR5 (3 days)-progressive ratio(PR) (1 day) schedule.

### TeNT-mediated synaptic silencing

4 weeks after the viral vector injection, rats were subjected to jugular catheterization and recovered for 1 week. Then rats were trained for heroin SA under FR1 schedule in context A for 10 days and extinction in context B for at least 7 days. 24 h after the last extinction training session, rats were re-introduced into context A for context-induced relapse test.

### Single-nucleus RNA seq

#### Tissue preparation

A total of 23 adult male rats randomly divided into three groups (Heroin-ABA, Heroin-ABB and Saline-ABB) were used for single-nuclei RNA-sequencing. Rats were deeply anesthetized by Zoletil, and brains were promptly dissected. Coronal slices containing the entire RSG were cut down with a thickness of 1 mm and the RSG tissues were microdissected cautiously under a stereo-microscope. The RSG tissues from each experimental group were pooled into the same reaction. Samples were then stored at –80℃ until subsequent experiments.

#### Single nucleus isolation and sequencing

The nuclei isolation and sequencing were performed by OE Biotech Co. Ltd (Shanghai, China). Preparation of nuclei suspension was conducted under ice cold conditions. In brief, frozen tissue samples were minced in 1ml of cell lysate and incubated for 7 minutes. Following the lysis verification by staining with trypan blue, 1ml wash buffer was added and the suspension was filtered by 40-μm cell strainers. The filtrate was transferred to centrifuge tubes and the cell strainer was washed with wash buffer and the rinse solution was combined with the nuclei filtrate. Then the suspension was centrifuged at 4℃, 500g for 5 minutes. The cell nuclei were resuspended in 5ml PBS with 1% BSA, then washed and centrifuged under the same conditions. The nuclei were again resuspended in 100 µl PBS with 1% BSA. After Trypan blue staining, the number of cell nuclei was counted by microscopy. Next, nuclei were diluted to 700-1200/µl using PBS with 1% BSA and the cDNA was amplified using the Next GEM Single Cell 3ʹ Reagent Kits v3.1. DNA library construction was performed followed by sequencing using the PE150 sequencing mode on the Illumina Nova 6000 platform.

#### Data processing and analysis

The official software CellRanger from 10×Genomics was employed for the preliminary quality control and reference genome alignment. Then the Seurat package facilitated the further filtration of low-quality cells. The high-quality cells that were preserved were defined as: gene count and Unique Molecular Identifier (UMI) count = median ± 2 × MAD with the mitochondrial UMI reads < 20%. Doublets were removed by the DoubletFinder package.

The ‘FindVariableGenes’ function in Seurat package facilitated the screen of highly variable genes which was used to conduct PCA dimensionality reduction analysis. The two-dimensional visualization was acquired by the Uniform Manifold Approximation and Projection (UMAP).

Marker genes identification was performed using the ‘FindAllMarkers’ function in Seurat package, which were then used to define the cell types of each cell cluster. Based on the public reference dataset, SingleR package facilitated the cell type definition via calculation of the correlation between the gene expression profile and the reference dataset. The cell type of each cluster was assigned according to the correlation with the reference dataset.

To further investigate the neuronal transcriptome, the identified neuronal cluster was then subjected to dimensionality reduction, re-clustering, differential gene expression analysis and KEGG and GO enrichment analysis. To investigate the heroin-addiction-related transcriptional alteration, differentially expressed IEGs from each neuronal cluster of Heroin-ABA *vs* Heroin-ABB, Heroin-ABB *vs* Saline-ABB were analyzed.

### Stereo-seq for spatial transcriptome analysis

#### Tissue preparation

After 90-min context-induced relapse test, rat was deeply anesthetized by Zoletil and brain was gently collected, embedded in OCT and frozen by liquid nitrogen followed by transferring to –80 ℃ freezer. Brain slices with 10 μm thickness were obtained using a cryostat microtome (Leica CM3050S). Then the slices were mounted onto the Stereo-seq chip and incubated at 37℃ for 3 min followed by methanol fixation for 40 min at –20℃. Slices staining was conducted using nucleic acid dye and digitally scanned via Stereo OR 100 microscope. The spatial transcriptome sequencing was conducted by Novogene Co. Ltd (Beijing, China).

#### In situ reverse transcription

Brain slices on the chip were washed in SSC buffer with RNase inhibitor followed by permeabilization by 0.1% pepsin and incubating at 37℃ for 5 min. The slices were again washed in SSC buffer. RNA released from the slices was captured by DNA Nano Ball (DNB) pre-designed on the chip and the reverse transcription was performed at 42℃ overnight using SuperScript II. The chip with cDNAs was then treated with Prepare cDNA Release Mix overnight at 55℃. cDNAs were subsequently purified using the DNA Clean Beads and amplified with KAPA HiFi Hotstart Ready Mix.

#### Library construction and sequencing

First, the QubitTM dsDNA Assay Kit was used to determine the concentrations of the amplified cDNA. Then 20 ng DNA were fragmented at 55℃ for 10 min using Tn5 transposase and were subsequently amplified in the mixed solution as follows: 25 ml fragmented DNA, 1×KAPA HiFi Hotstart Ready Mix and 0.3 mM Stereo-seq-Library-F/R primer with supplement of nuclease-free water to 100 ml. The reaction was then succeeded by 2 consecutive heating cycles followed by purification by the AMPure XP Beads and the sequencing was conducted using the MGI DNBSEQ-Tx sequencer.

#### Data processing and analysis

Data containing the Coordinate Identifier (CID) and Multiple Identification Domain (MID) sequences from the Stereo-seq chip and cDNA sequences was produced by MGI DNBSEQ-Tx sequencer. The CID sequences, carrying the absolute spatial information on the Stereo-seq chip, were mapped to the corresponding coordinates of the chip. MID, carrying the spatial information of the captured RNA, and CID sequences were added to each read which were next aligned to the reference genome. Reads with Mapping Quality > 10 were annotated to the corresponding genes and counted. UMIs with the same sequences of CID were combined. These information were finally integrated to achieve a spatial matrix.

The cell segregation was conducted based on the image of nucleic acid staining and the spatial matrix was then divided into non-overlapping cell bins for further dimension reduction, clustering, cell type definition and differentially expressed gene analysis. The UMAP and spatial distribution visualization and the following quantitative analysis were processed on DCS Cloud (BGI Tech, China).

### Tissue clearing

#### Tissue preparation

Rats were deeply anesthetized by Zoletil followed by transcardial perfusion with sterile 1×PBS and 4% paraformaldehyde (PFA). Brains were then collected and immersed in 4% PFA for 24 h at 4℃, and subsequently stored in PBS at 4℃ until use.

#### SHIELD for brain clearing

Brains were first incubated in SHIELD OFF solution (20ml SHIELD-Epoxy, 10ml SHIELD Buffer and 10ml ddH2O) at 4℃ for 6 days with light shaking. Then samples were transferred to 40ml SHIELD ON solution and incubated for 24 h at 37 ℃. Subsequently, brains were rinsed in Delipidation Buffer for 24 hours at 45°C followed by active delipidation under 40V, 1250 mA and 42℃ conditions for 6 days. The refraction index matching was then conducted by incubating the brains in EasyIndex solution (RI=1.52) until transparent. The 3D imaging was achieved using Light Sheet Fluorescence Microscopy.

### Fluorescence in situ hybridization

#### Tissue preparation

After the 90-min behavioural test, rats were deeply anesthetized by Zoletil followed by transcardial perfusion with the DEPC-treated 1×PBS and 4% paraformaldehyde (PFA). Brains were then collected and immersed in 4% PFA for 4 h at 4℃, and subsequently immersed into 30% sucrose solution for 72 h at 4℃. Brain slices with 20 μm thickness were obtained using a cryostat microtome and mounted onto SuperFrost Plus microscope slides. Brain slices were stored at –80℃ until use.

#### mRNA detection

mRNA expression detection in RSG, ZI, AD, RSA, ACC, M2, Cla, HDB and Sub was conducted using RNAscope multiplex fluorescent reagent kit v2 assay. Briefly, brain slices were first dehydrated and treated with tissue antigen retrieval reagent, then incubated with RNAScope protease III at 40 °C for 30 min. Then the mRNA probes targeting *Vglut1* (cat.# 317001), *Vglut2* (cat. # 317011-C2), *Vgat* (cat. # 424541), *Cckbr* (cat. # 312001-C2), *Cck* (cat. # 532851-C2), *mCherry* (cat. # 431201-C3), *EGFP* (cat. # 538851-C3), *Fos* (cat. # 403591-C3) were added to the slices and incubated at 40 °C for 2 h. The signal amplification was conducted to obtain high detection qualities.

### Immunohistochemistry

#### Tissue preparation

After the 90-min behavioural test, rats were deeply anesthetized by Zoletil followed by transcardial perfusion with 1×PBS and 4% paraformaldehyde (PFA). Brains were then collected and immersed in 4% PFA at 4℃ for 6 h, and subsequently dehydrated in 30% sucrose solution for 72 h at 4℃. Brain slices with 40 μm thickness were obtained using a cryostat microtome and immersed in 1×PBS at 4℃ until use. The immunostaining experiment was conducted within 24 h after sectioning.

#### Protein immunostaining

Free-floating cryo-sections were first washed in 1×PBS 5 min for 3 times. Then the slices were blocked in 3% normal goat serum solution with 0.3% Triton for 1.5 h followed by incubation with primary antibodies overnight at 4℃. Then the slices were washed in 1×PBS 5 min for 3 times and incubated in secondary antibodies for 2 h. Zeiss 710 laser confocal microscope was used for signal detection and imaging. The quantitative analysis was conducted by ImageJ software. Primary antibodies: rabbit anti-c-fos (1:1000, Synaptic systems), mouse anti-c-fos (1:300, abcam), mouse anti-CaMKIIα (1:300, abcam), rabbit anti-Cckbr (1:200, Alomone labs), mouse anti-Gad (1:200, abcam), rabbit anti-TH (1:1000, millipore). Secondary antibodies: goat anti-rabbit (Alexa Fluor 488), goat anti-mouse (Alexa Fluor 555), goat anti-rabbit (Alexa Fluor 647).

### Behavioral assays

#### Heroin self-administration

Heroin self-administration (heroin SA) training was conducted as previously described^99^ with minor modification. Briefly, rats with jugular catheterization were first introduced into context A and allowed 30-min free exploration for 3 consecutive days to acclimate to the environment. Then the infusion pump was connected to the intravenous silicone tube followed by the start of behavioral training software. For fixed ratio 1 (FR1) training schedule, rats were trained for 2 h every session for 10 consecutive days. For evaluation of the effect of caspase-mediated apoptosis, rats were trained under the following schedule: 3-day FR1, 3-day FR2, 3-day FR5 and 1-day progressive ratio (PR) program. The number of responses required by the next drug infusion under PR schedule was determined by the equation: number of responses = (5 × e ^(0.2 × infusion number)^)−5.

#### ABA renewal context-induced relapse

Briefly, rats undergoing 10-day heroin SA training in context A were introduced to context B for at least 7-day extinction training (2 h per session), during which the active poke behavior no longer resulted in drug delivery but still triggered a light-sound cue. 24 h after the last extinction training session, rats were re-introduced into context A for a 90-min relapse test without drug infusion.

#### Water licking and chocolate consumption

Rats were housed individually in their homecage for 3 weeks after stereotaxic surgery with ad libitum water and food. One day before the experiment, rats were allowed to acclimate to the chamber for 30 min. Then rats were fasted or water deprived for 12 h, then rats were introduced into the testing chamber with water or chocolate provided multiple times. Fiber photometry recording began as the rats got into the chamber.

#### Open field test

Rats with clozapine injection i.p. 30 min early were placed into the center of a cubic chamber (50 cm × 50 cm × 60 cm). A micro-camera was located directly over the chamber and was used to monitor the locomotor activity of each rat for 10 min. The data of locomotion was analyzed by JL Behv software (JiLiang Software Technology Co., Ltd, Shanghai, China).

## QUANTIFICATION AND STATISTICAL ANALYSIS

All data are shown as the mean ± SEM, unless otherwise stated. Statistical analyses were performed using R and PRISM 9.5.1 Software. Before the difference analysis, all data were tested for normal distribution. The parametric or non-parametric test was used according to whether the data were Gaussian distributions or not. The methods for difference analysis were stated in the corresponding figure legends. The ‘n’ in the data description indicates the number of rats or cells. Results were regarded as statistically significant with *p* < 0.05.

**Figure S1.**
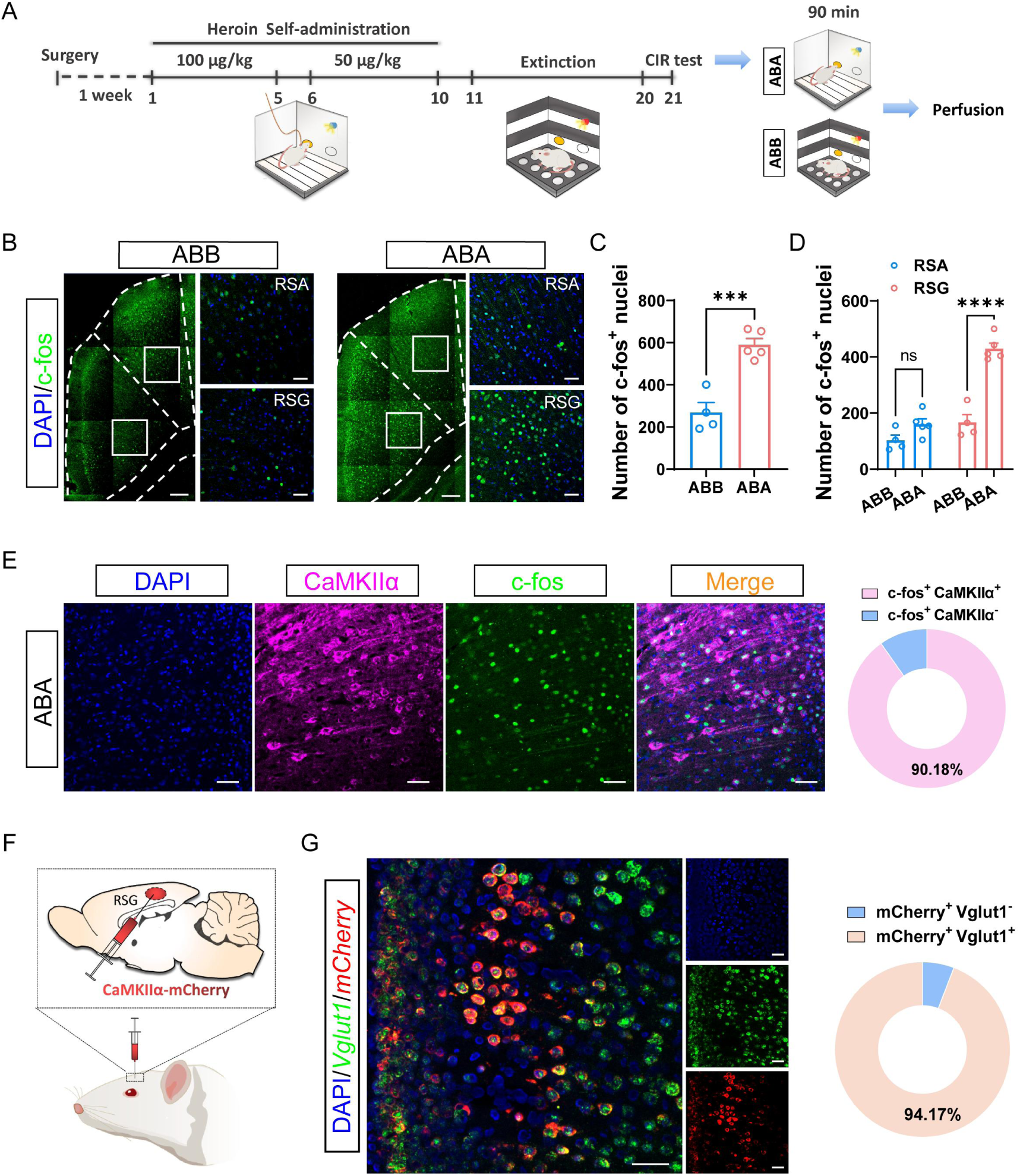
Activity of RSG glutamatergic neurons during context-induced relapse and validation of the virus specificity, related to figure 1. **(A)** Schematic showing the training schedule for heroin SA, extinction and context-induced relapse and the perfusion time point for immunostaining. **(B)** Representative images of the c-fos expression in RSC of rats in ABB or ABA groups. Scale bars, 200 μm and 50 μm. **(C)** Number of c-fos-positive nuclei in RSC of rats in ABB group (*n* = 4) *vs* ABA group (*n* = 5). Unpaired t test, \*\*\**p* < 0.001. **(D)** Number of c-fos-positive nuclei in RSG and RSA of rats in ABB group (*n* = 4) *vs* ABA group (*n* = 5). Two-way ANOVA (F_(1,14)_ = 56.14, *p* < 0.0001) followed by Sidak’s post hoc test, \*\*\*\**p* < 0.0001, ns, no significant difference. **(E)** Left: representative images showing the co-localization of c-fos and CaMKIIα in RSG after context-induced relapse test. Scale bars, 50 μm. Right: percentage of CaMKIIα-positive and negative cells in c-fos^+^ nuclei (*n* = 3). **(F)** Schematic showing the viral strategy for selective labeling of RSG glutamatergic neurons by AAVs carrying CaMKIIα promoter. **(G)** Left: representative images showing the co-localization of *Vglut1* and *mCherry* in RSG. Scale bars, 50 μm. Right: percentage of Vglut1-positive and negative cells in mCherry-labeled neurons in RSG (*n* = 3).

**Figure S2.**
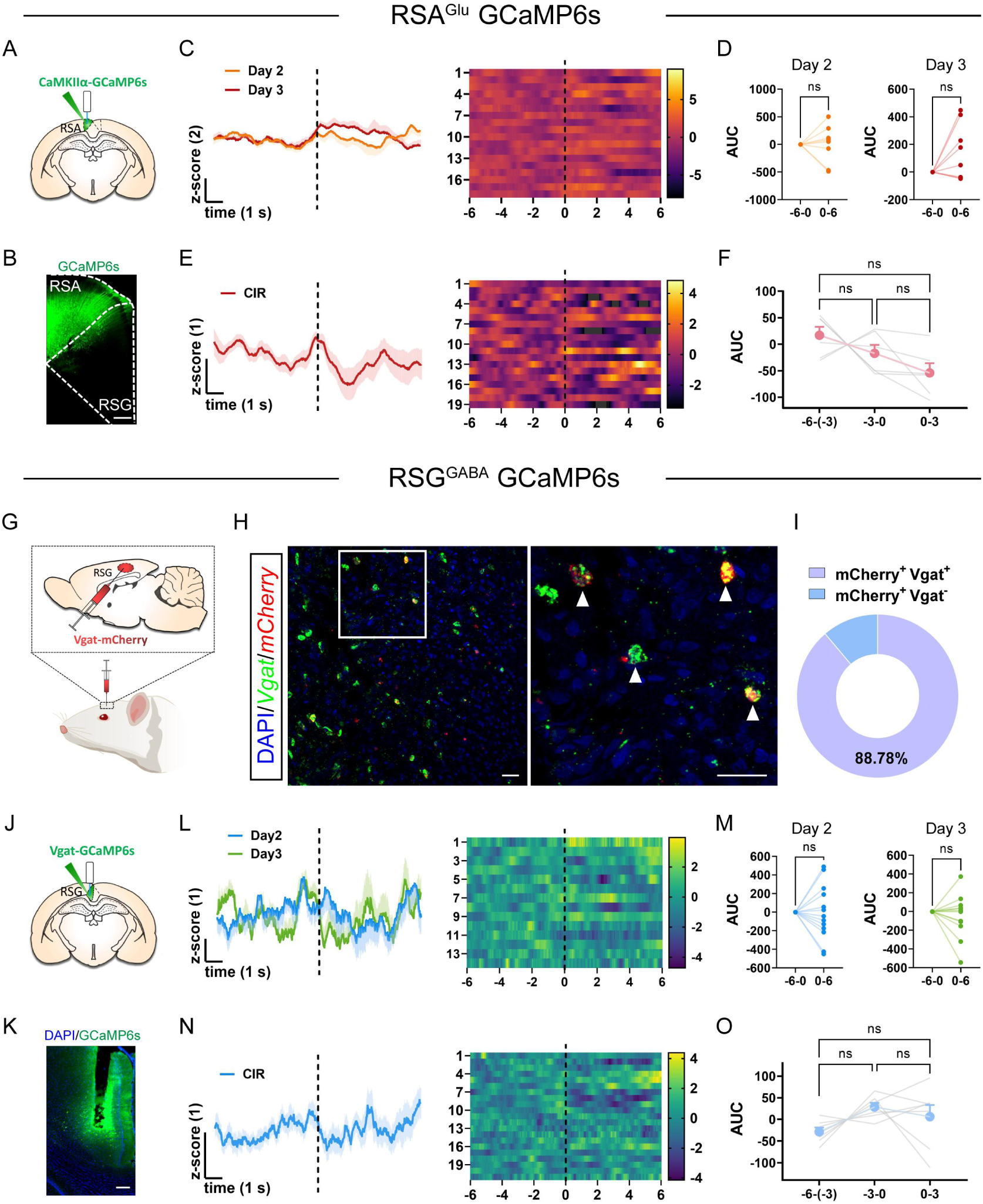
Activity of RSA glutamatergic neurons and RSG GABAergic neurons during heroin SA training and context-induced relapse and validation of the virus specificity, related to figure 1. **(A)** Schematic showing the viral strategy and fiber implantation for Ca^2+^ signal recording of RSA glutamatergic neurons. **(B)** Representative image of the expression of GCaMP6s in RSA. Scale bars, 200 μm. **(C)** Average Ca^2+^ signals and heatmaps of RSA^Glu^ neurons during day 2-3 of heroin SA training. Vertical dashed line indicates the onset of active poke behaviors. **(D)** Area under curve of the RSA^Glu^ Ca^2+^ signals during day 2 (*n* = 8) and day 3 (*n* = 8) of heroin SA training. Paired t test, ns, no significant difference. **(E)** Average Ca^2+^ signals and heatmaps of RSA^Glu^ neurons during context-induced relapse test. Vertical dashed line indicates the onset of active poke behaviors. **(F)** Area under curve of the RSA^Glu^ Ca^2+^ signals during context-induced relapse test (*n* = 6). RM One-way ANOVA (F_(1.508, 7.538)_ = 3.453, *p* = 0.0936) followed by Tukey’s post hoc test, ns, no significant difference. **(G)** Schematic showing the viral strategy for selective labeling of RSG GABAergic neurons by AAVs carrying Vgat promoter. **(H)** Representative images showing the co-localization of *Vgat* and *mCherry* in RSG. Scale bars, 50 μm. **(I)** Percentage of Vgat-positive and negative cells in mCherry-labeled neurons in RSG (*n* = 3). **(J)** Schematic showing the viral strategy and fiber implantation for Ca^2+^ signal recording of RSG GABAergic neurons. **(K)** Representative image of the expression of GCaMP6s in RSG. Scale bars, 200 μm. **(L)** Average Ca^2+^ signals and heatmaps of RSG GABAergic neurons during day 2-3 of heroin SA training. Vertical dashed line indicates the onset of active poke behaviors. **(M)** Area under curve of the RSG^GABA^ neurons Ca^2+^ signals during day 2 (*n* = 13) and day 3 (*n* = 12) of heroin SA training. Paired t test, ns, no significant difference. **(N)** Average Ca^2+^ signals and heatmaps of RSG^GABA^ neurons during context-induced relapse test. Vertical dashed line indicates the onset of active poke behaviors. **(O)** Area under curve of the RSG^GABA^ neurons Ca^2+^ signals during context-induced relapse test (*n* = 7). RM One-way ANOVA (F_(1.622, 9.733)_ = 2.482, *p* = 0.1398) followed by Tukey’s post hoc test, ns, no significant difference.

**Figure S3.**
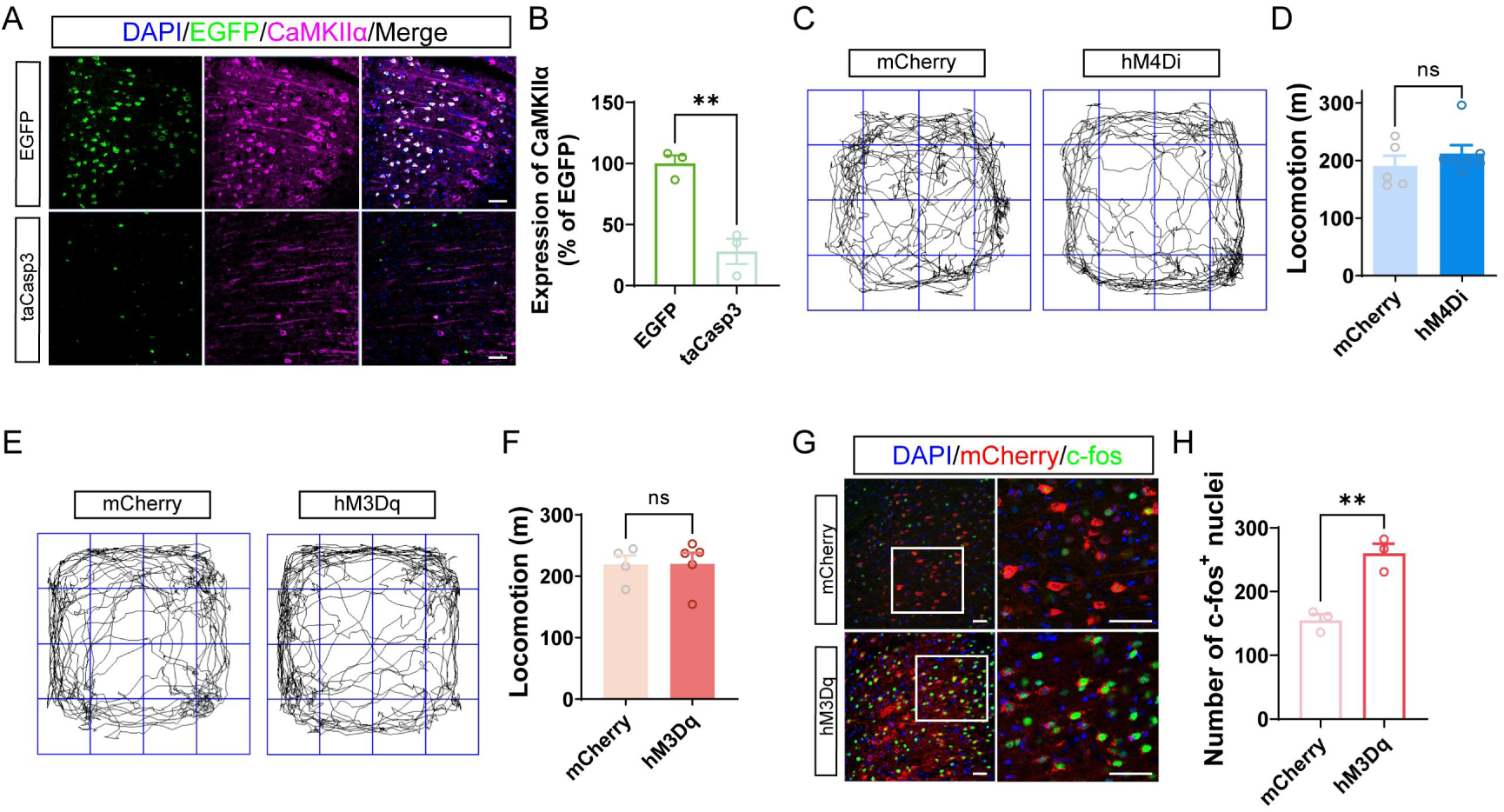
Validation of the apoptosis and chemogenetic activation of RSG^Glu^ neurons and the effect of chemogenetic manipulation on the locomotion of rats, related to figure 1. **(A)** Representative images showing the expression of CaMKIIα in RSG of rats in EGFP control and taCasp groups. Scale bars, 50 μm. **(B)** Normalized CaMKIIα expression in RSG of EGFP control group (*n* = 3) *vs* taCasp group (*n* = 3). Unpaired t test, \*\**p* < 0.01. **(C)** Representative tracks of rats in mCherry control and hM4Di groups. **(D)** Quantification of locomotion following clozapine injection in mCherry-expressing (*n* = 5) and hM4Di-expressing rats (*n* = 7). Mann-Whitney test, ns, no significant difference. **(E)** Representative tracks of rats in mCherry control and hM3Dq groups. **(F)** Quantification of locomotion following clozapine injection in mCherry-expressing (*n* = 4) and hM3Dq-expressing rats (*n* = 5). Unpaired t test, ns, no significant difference. **(G)** Representative images showing the expression of c-fos in RSG of rats in mCherry control and hM3Dq groups. Scale bars, 50 μm. **(H)** Number of c-fos-positive nuclei in mCherry control group (*n* = 3) *vs* hM3Dq group (*n* = 3). Unpaired t test, \*\**p* < 0.01.

**Figure S4.**
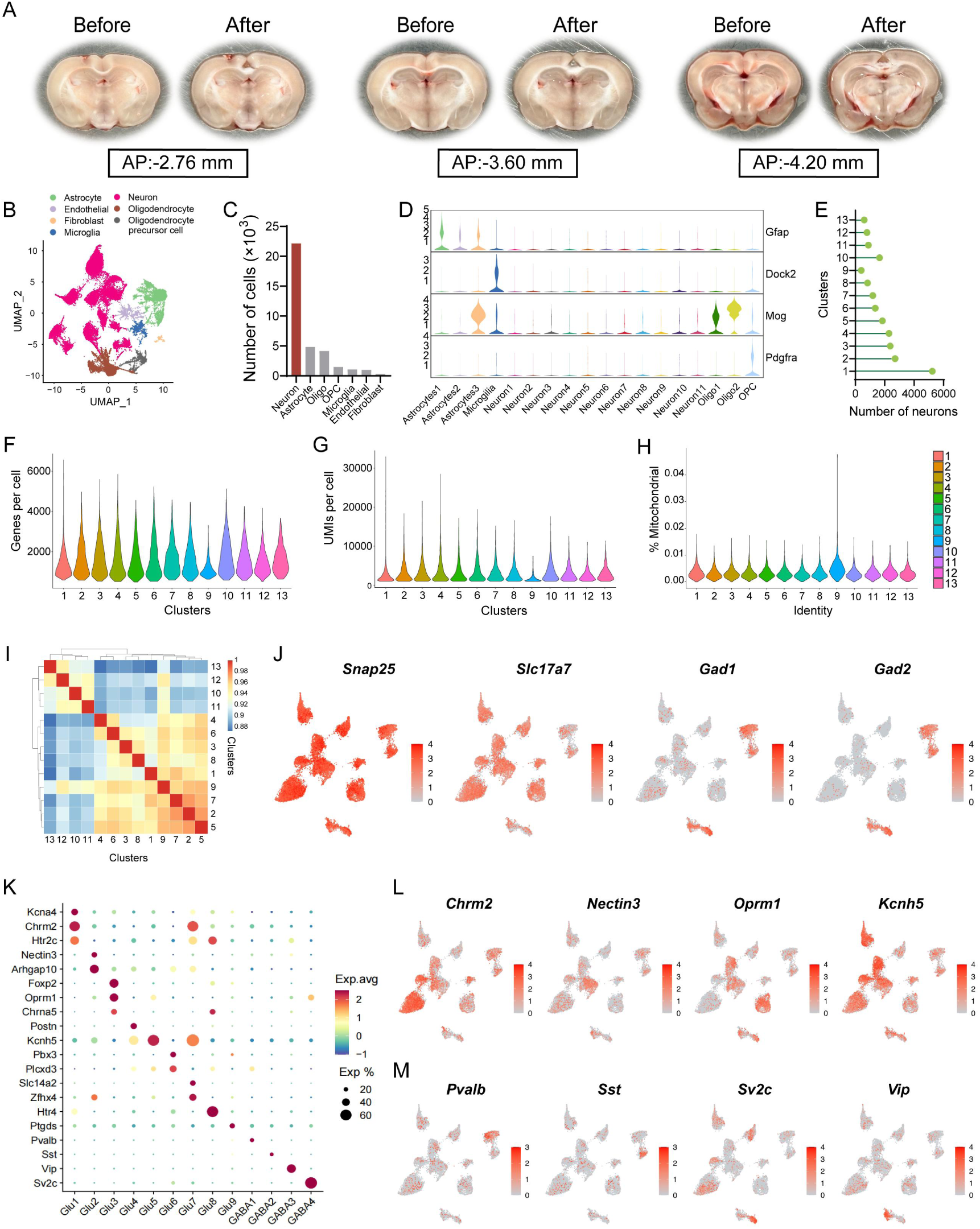
Additional information for snRNA-seq, related to figure 2. **(A)** Representative images showing the microdissection of the RSG in coronal slices. **(B)** UMAP visualization of 35019 cells of RSG. **(C)** Number of each type of cells in RSG. **(D)** Violin plots of marker genes for each type of cells in RSG. **(E)** Number of neurons in each neuronal cluster. **(F)** Number of genes per cell in each neuronal cluster. **(G)** Number of UMI per cell in each neuronal cluster. **(H)** Percentage of mitochondrial gene reads per cell in each neuronal cluster. **(I)** Co-efficiency heatmap of all neuronal clusters. **(J)** Expression of marker genes used to identify neurons, glutamatergic neurons and GABAergic neurons. **(K)** Dotplots showing the averaged expression levels (color) and the percentage of expressing neurons (size) of specific marker genes in each neuronal cluster. **(L)** UMAP visualization of the distribution of some marker genes of glutamatergic neuronal clusters. **(M)** UMAP visualization of the distribution of some marker genes of GABAergic neuronal clusters.

**Figure S5.**
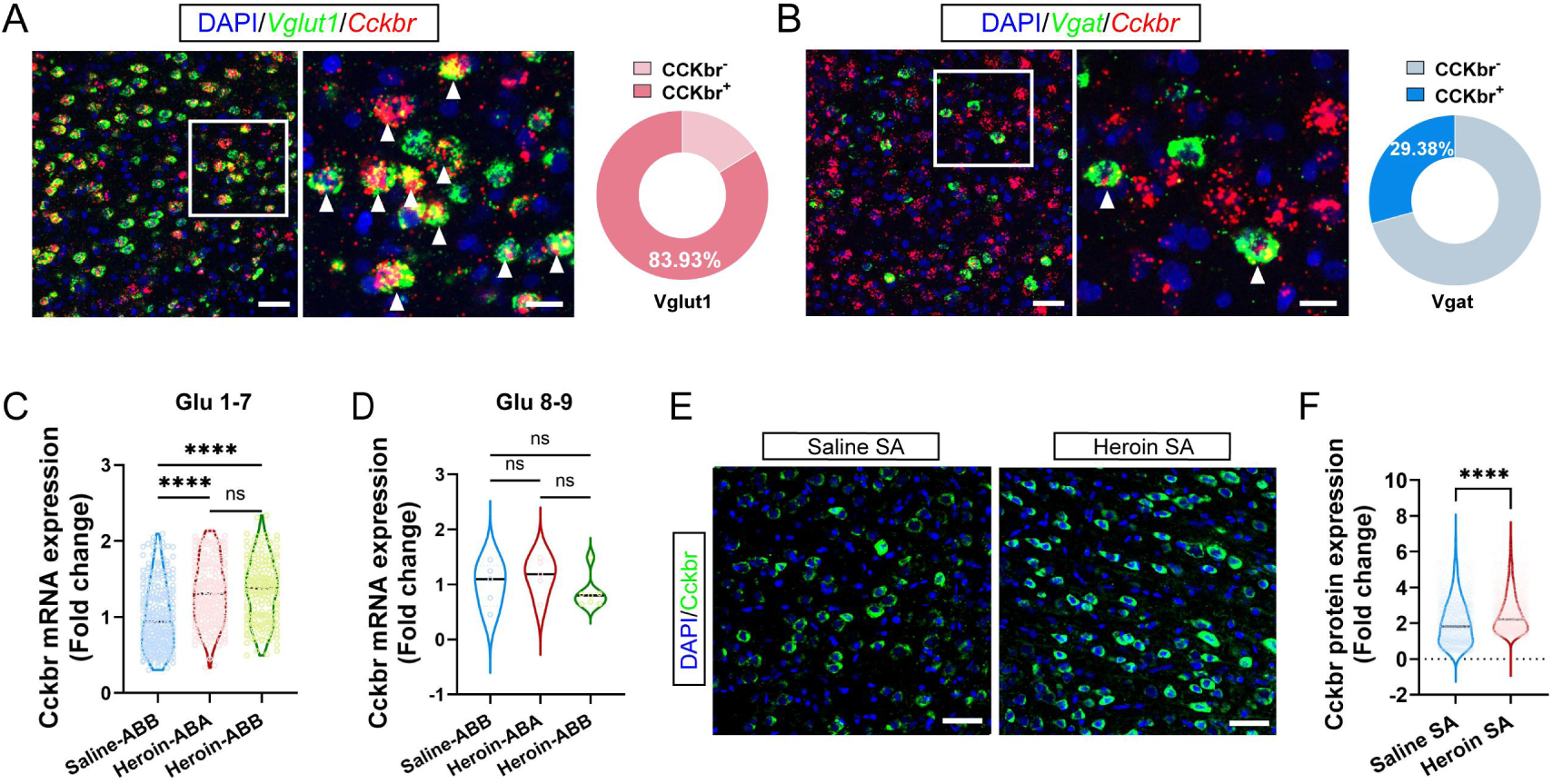
Validation of the dominant Cckbr expression in RSG^Vglut1^ neurons and the increased Cckbr expression in RSG induced by heroin SA training, related to figure 2. **(A)** Left: representative images showing the expression of *Vglut1* and *Cckbr* in RSG. Right: percentage of Cckbr^+^ and Cckbr^-^ cells in Vglut1^+^ neurons (*n* = 5). Scale bars, 50 μm and 20 μm. **(B)** Left: representative images showing the expression of *Vgat* and *Cckbr* in RSG. Right: percentage of Cckbr^+^ and Cckbr^-^ cells in Vgat^+^ neurons (*n* = 5). Scale bars, 50 μm and 20 μm. **(C-D)** Average expression level of *Cckbr* mRNA of Cckbr-positive neurons in clusters Glu 1-7 (C) and Glu 8-9 (D) in Saline-ABB, Heroin-ABA and Heroin-ABB groups from snRNA-seq data. Kruskal-Wallis test followed by Dunn’s test, \*\*\*\**p* < 0.0001, ns, no significant difference. **(E)** Representative images showing the expression of Cckbr protein in RSG of Saline SA and Heroin SA rats. Scale bars, 50 μm. **(F)** Average expression level of Cckbr protein in RSG of Saline SA (*n* = 3) *vs* Heroin SA (*n* = 3) rats. Mann-Whitney test, \*\*\*\**p* < 0.0001.

**Figure S6.**
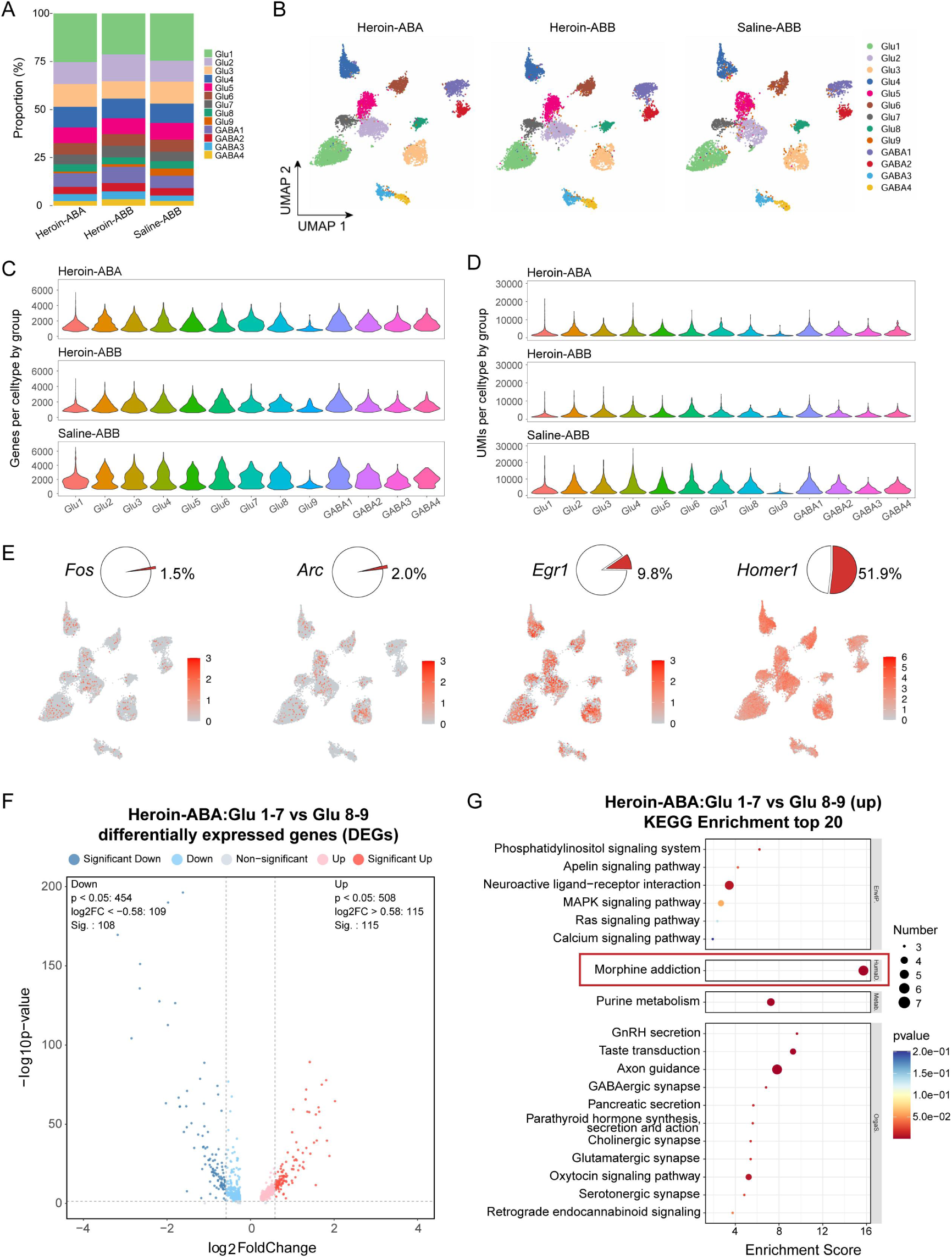
Additional information for snRNA-seq, related to figure 2. **(A)** Proportions of each neuronal cluster in Heroin-ABA, Heroin-ABB and Saline-ABB groups. **(B)** UMAP visualization of neuronal clusters in Heroin-ABA, Heroin-ABB and Saline-ABB groups. **(C-D)** Number of genes (C) and UMIs (D) per cell in each neuronal cluster of Heroin-ABA, Heroin-ABB and Saline-ABB groups. **(E)** Expression of some representative IEGs in neuronal clusters visualized by UMAP. **(F)** Volcano plot showing the differentially expressed genes (DEGs) between Glu1-7 and Glu 8-9 in Heroin-ABA group. **(G)** KEGG analysis of the DEGs between Glu1-7 and Glu 8-9 in Heroin-ABA group. The Morphine addiction pathway in the red box is enriched in Human Disease (HumaD.).

**Figure S7.**
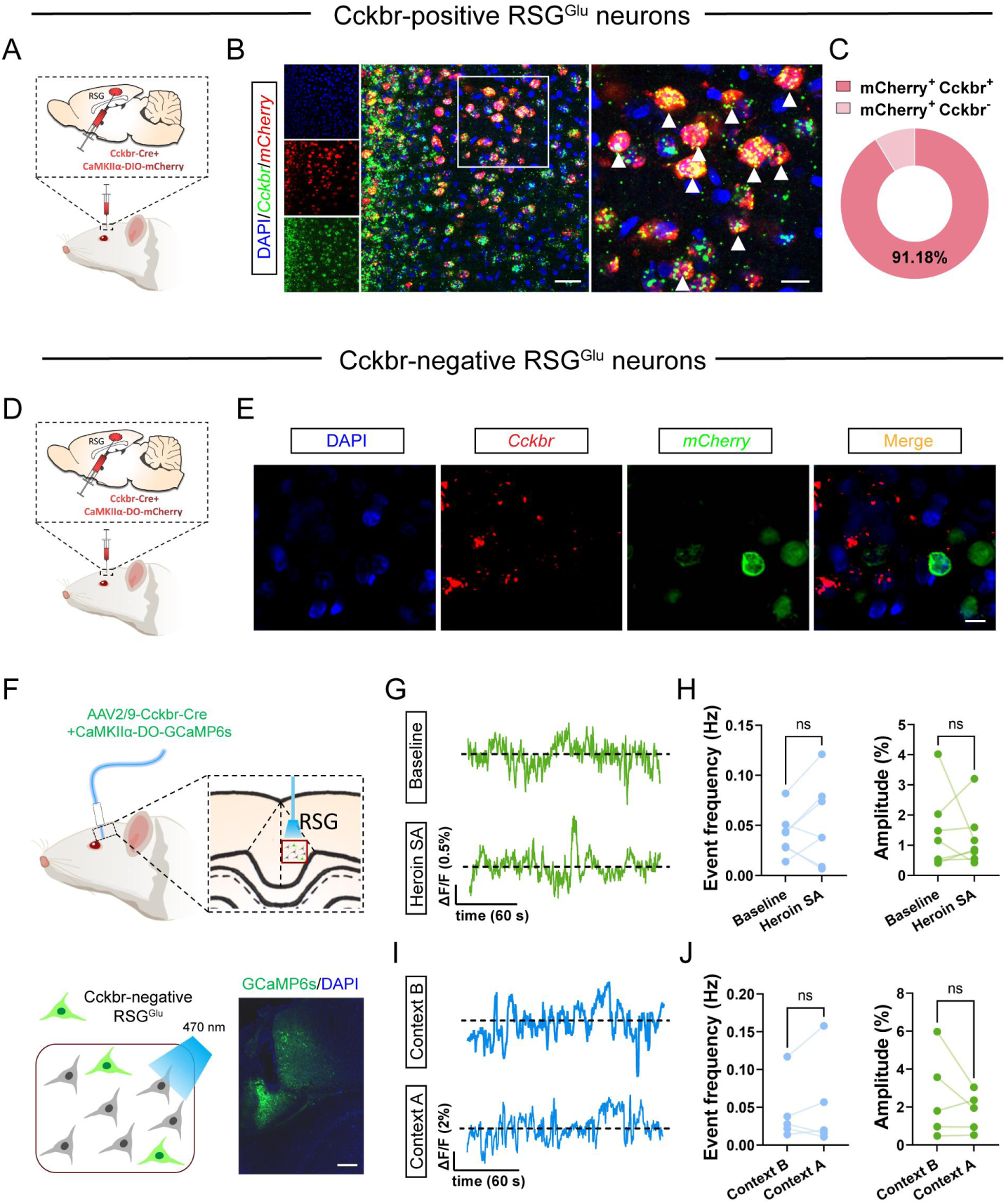
Validation of the virus specificity and the Ca^2+^ activity of Cckbr negative glutamatergic neurons in RSG, related to figure 3. **(A)** Schematic showing the viral strategy for selective labeling of RSG Cckbr-positive glutamatergic neurons. **(B)** Representative images showing the co-localization of *mCherry* and *Cckbr* in RSG. Scale bars, 50 μm and 20 μm. **(C)** Percentage of Cckbr-positive and negative cells in mCherry^+^ neurons (*n* = 3). **(D)** Schematic showing the viral strategy for selective labeling of RSG Cckbr-negative glutamatergic neurons. **(E)** Representative images showing the separate distribution of *mCherry* and *Cckbr* in RSG. Scale bars, 10 μm. **(F)** Top: Schematic showing the viral strategy and fiber implantation for fiber photometry recording of Cckbr-negative glutamatergic neurons in RSG. Bottom: representative image showing the expression of GCaMP6s in RSG. Scale bars, 200 μm. **(G-H)** Representative traces (G) and the event frequency and average amplitude (H) of the tonic Ca^2+^ signal of RSG Cckbr-negative glutamatergic neurons during baseline recording *vs* heroin SA training (*n* = 7). Paired t test, ns, no significant difference. **(I-J)** Representative traces (I) and the event frequency and average amplitude (J) of the tonic Ca^2+^ signal of RSG Cckbr-negative glutamatergic neurons during extinction in context B *vs* relapse test in context A.Wilcoxon matched-pairs signed rank test for event frequency and Paired t test for amplitude, ns, no significant difference.

**Figure S8.**
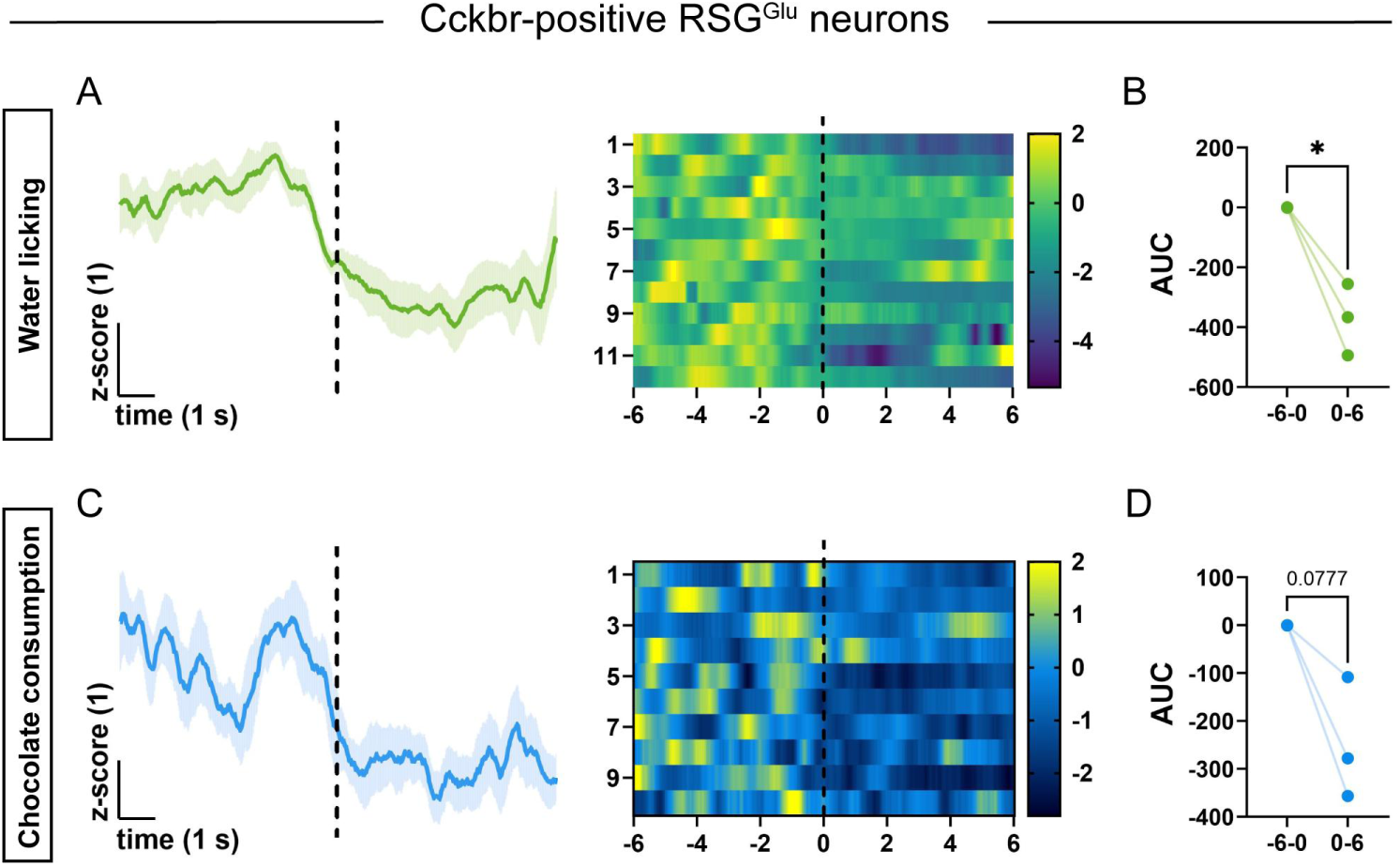
Ca^2+^ activity of Cckbr positive glutamatergic neurons in obtaining natural reward, related to figure 3. **(A-D)** Average Ca^2+^ signals, heatmaps and area under curve of the RSG Cckbr-positive glutamatergic neurons during water licking (*n* = 3) and chocolate consumption (*n* = 3). Paired t test, \**p* < 0.05. Vertical dashed line indicates the onset of water licking and chocolate consumption.

**Figure S9.**
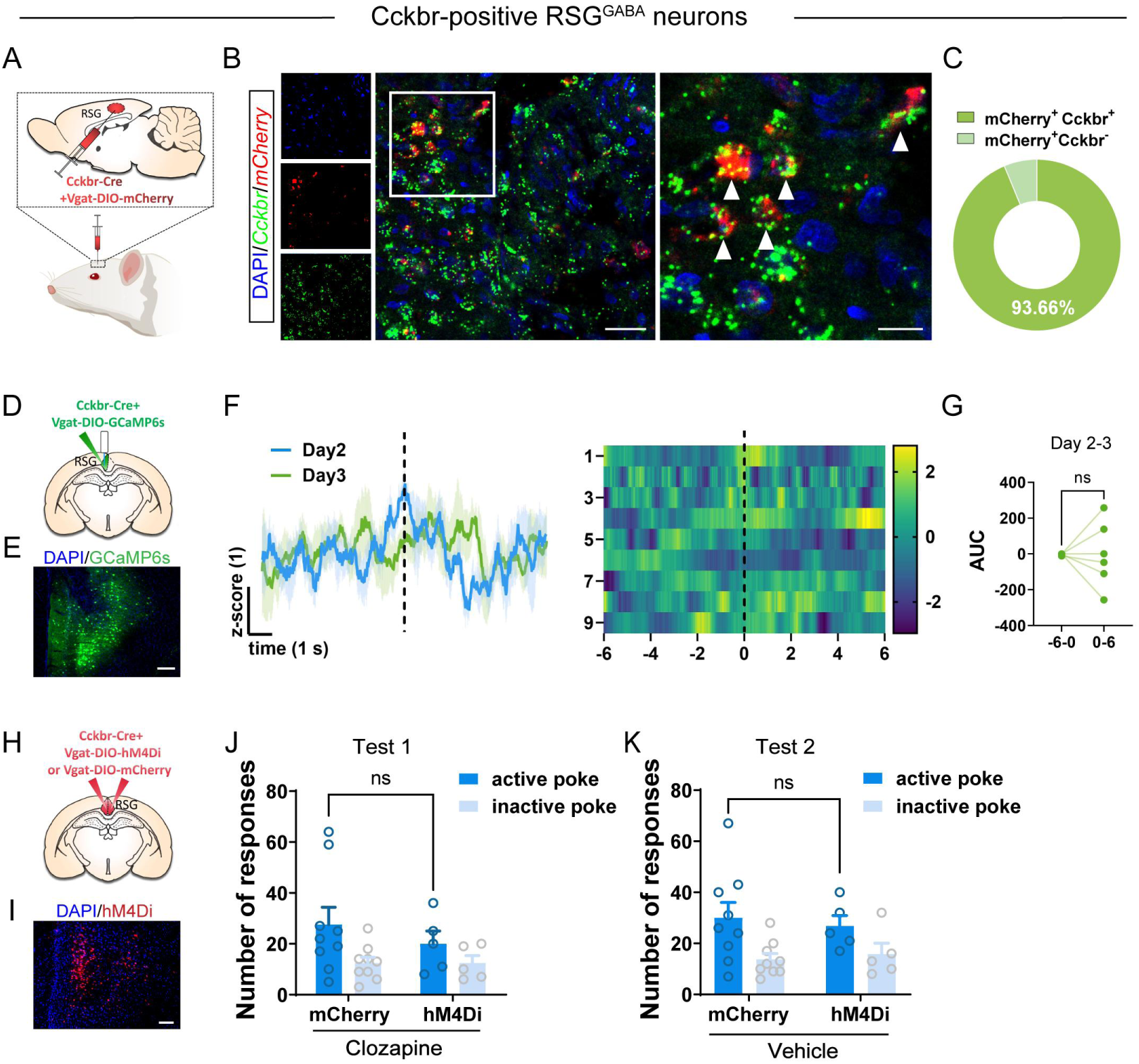
Activity of RSG^GABA-Cckbr^ neurons during heroin SA training and chemogenetic manipulation of RSG^GABA-Cckbr^ neurons during context-induced relapse, related to figure 3. **(A)** Schematic showing the viral strategy for selective labeling of Cckbr-positive GABAergic neurons in RSG. **(B)** Representative images showing the co-localization of *mCherry* and *Cckbr* in RSG. Scale bars, 50 μm and 20 μm.. **(C)** Percentage of Cckbr-positive and negative cells in mCherry^+^ neurons (*n* = 3). **(D)** Schematic showing the viral strategy and fiber implantation for selective recording of RSG Cckbr-positive GABAergic neurons. **(E)** Representative image showing the expression of GCaMP6s in RSG. Scale bars, 200 μm. **(F)** Average Ca^2+^ signals and heatmaps of RSG Cckbr-positive GABAergic neurons during day 2 and day 3 heroin SA training sessions. Vertical dashed line indicates the onset of active poke behaviors. **(G)** Area under curve of the Ca^2+^ signals of the Cckbr-positive GABAergic neurons in RSG during day 2-3 of heroin SA training (*n* = 6). Wilcoxon matched-pairs signed rank test, ns, no significant difference. **(H)** Schematic showing the viral strategy for selectively inhibition of RSG Cckbr-positive GABAergic neurons. **(I)** Representative image showing the expression of hM4Di in RSG. Scale bars, 200 μm. **(J-K)** Number of responses of rats in mCherry control group (*n* = 9) *vs* hM4Di group (*n* = 5) during relapse test 1 with clozapine injection or relapse test 2 with vehicle injection (i.p.). For test 1, Two-way ANOVA (F_(1,24)_ = 0.5048, *p* = 0.4843) followed by Sidak’s post hoc test; For test 2, Two-way ANOVA (F_(1,24)_ = 0.01443, *p* = 0.9054) followed by Sidak’s post hoc test, ns, no significant difference.

**Figure S10.**
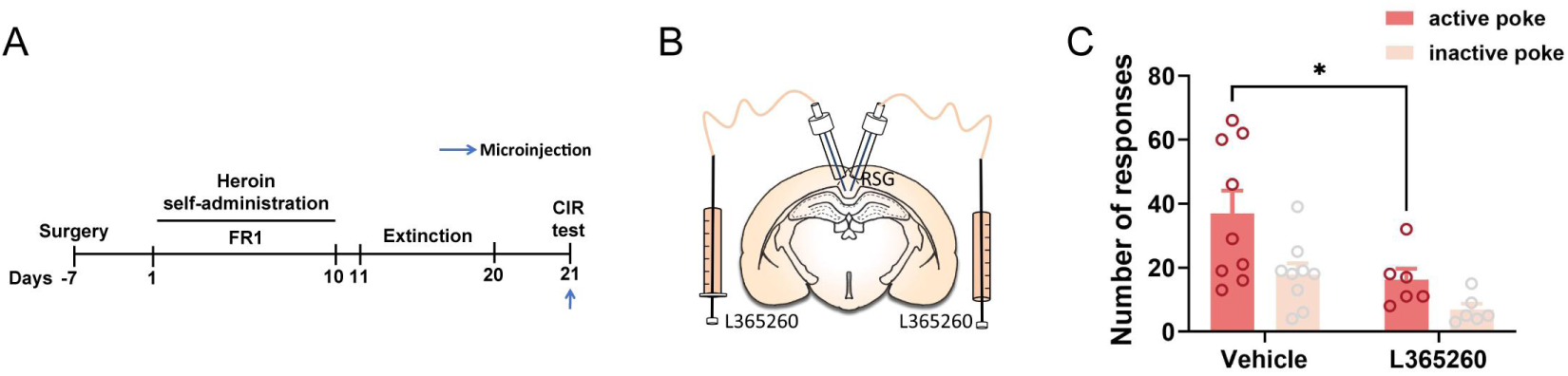
Effect of Cckbr antagonism on the context-induced relapse behavior, related to figure 4. **(A)** Schematic showing the training schedule and the time point for microinjection *in vivo*. **(B)** Schematic showing the strategy for bilateral RSG microinjection of L365260. **(C)** Number of responses of rats with vehicle (*n* = 9) *vs* L365260 (*n* = 6) injection into RSG. Two-way ANOVA (F_(1,26)_ = 9.323, *p* < 0.01) followed by Sidak’s post hoc test, \**p* < 0.05.

**Figure S11.**
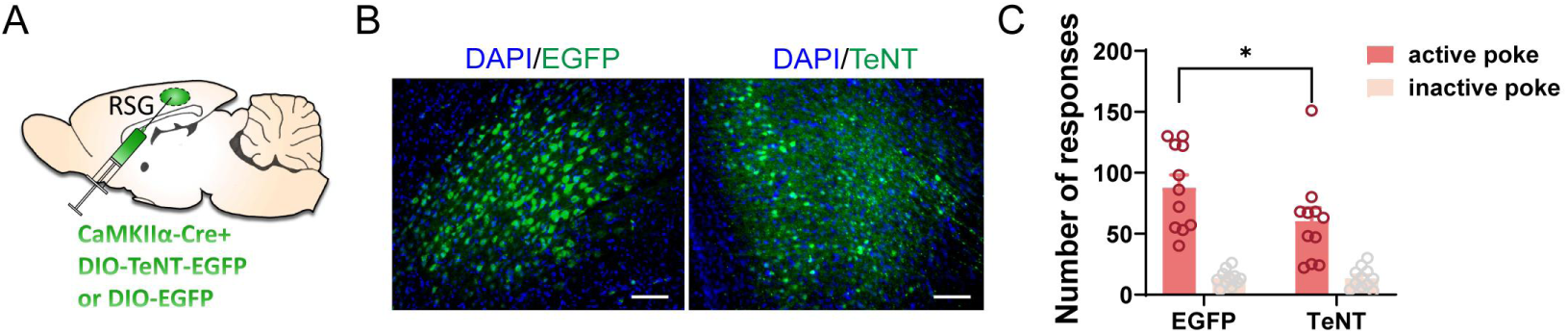
Effect of TeNT-mediated blocking of RSG^Glu^ synaptic transmission on the context-induced relapse behavior, related to figure 5. **(A)** Schematic showing the viral strategy for selective expression of TeNT in RSG glutamatergic neurons. **(B)** Representative images showing the expression of EGFP and TeNT in RSG. Scale bars, 100 μm. **(C)** Number of responses of rats in EGFP control group (*n* = 11) and TeNT group (*n* = 11) during context-induced relapse. Two-way ANOVA (F_(1,40)_ = 3.213, *p* = 0.0806) followed by Sidak’s post hoc test, \**p* < 0.05.

**Figure S12.**
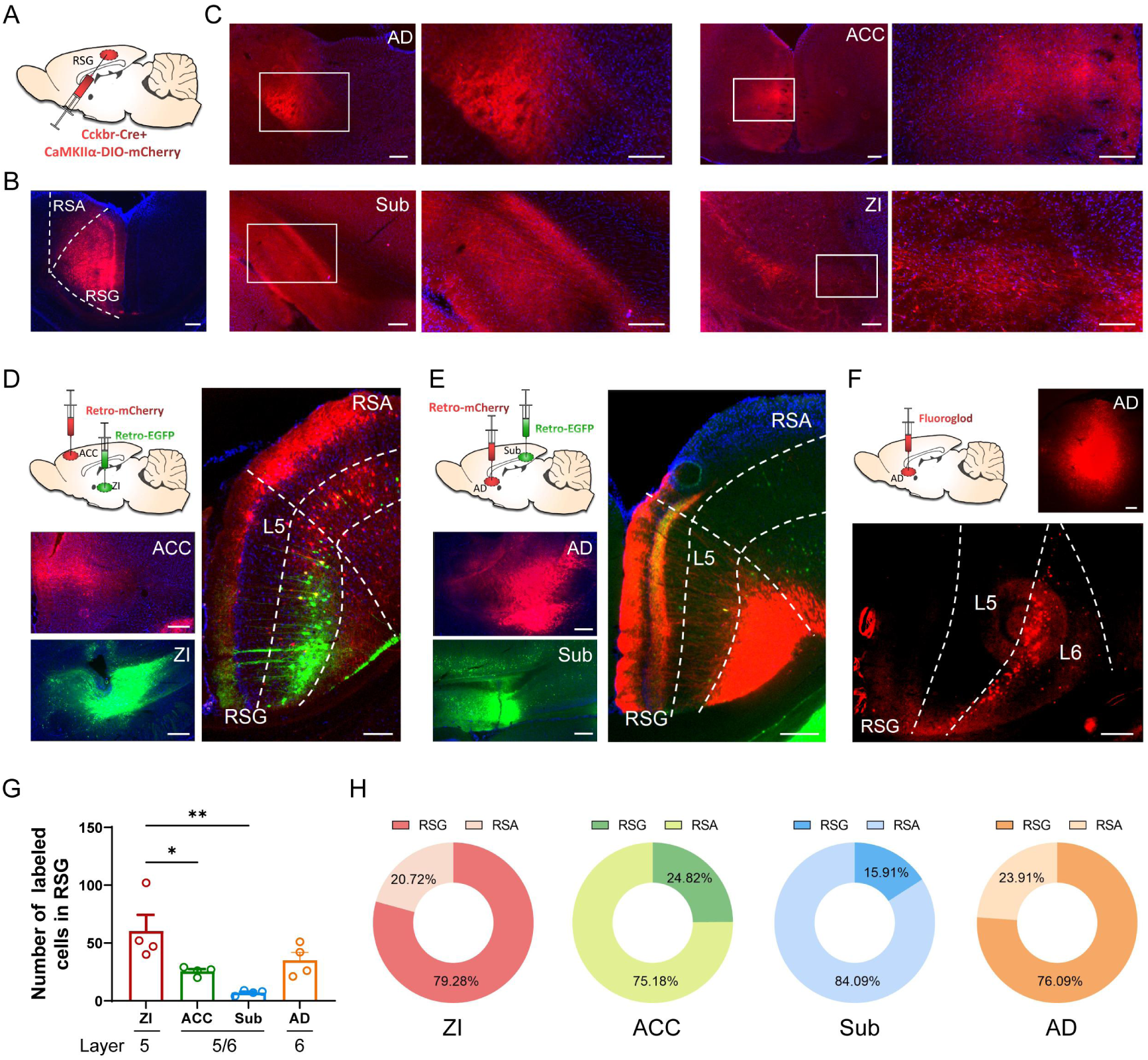
Projection profile of RSG^Glu-Cckbr^ neurons, related to figure 5. **(A)** Schematic showing the viral strategy for selective labeling of RSG^Glu-Cckbr^ neurons. **(B)** Representative image showing the expression of mCherry in RSG. Scale bars, 200 μm. **(C)** Axon terminal of the RSG^Glu-Cckbr^ neurons in different projection targets. Scale bars, 200 μm. AD: anterodorsal nucleus of thalamus, ACC: anterior cingulate cortex, Sub: subiculum, ZI: zona incerta. **(D)** Left: schematic showing the viral strategy for retrograde tracing from ZI and ACC (top) and representative images of the virus expression in ZI and ACC (bottom). Right: representative image showing the retrograde-labeled neurons in RSG. Scale bars, 200 μm. **(E)** Left: schematic showing the viral strategy for retrograde tracing from AD and Sub (top) and representative images of the virus expression in AD and Sub (bottom). Right: representative image showing the retrograde-labeled neurons in RSG. Scale bars, 200 μm. **(F)** Top: schematic showing the strategy for retrograde tracing from AD by fluorogold (left) and representative images of the fluorogold expression in AD (right). Bottom: representative image showing the retrograde-labeled neurons in RSG. Scale bars, 200 μm. **(G)** Number of retrograde-labeled neurons in RSG from different downstream targets (*n* = 4). One-way ANOVA (F_(3, 12)_ = 7.765, *p* < 0.01) followed by Tukey’s post hoc test, \**p* < 0.05, \*\**p* < 0.01. **(H)** Proportion of retrograde-labeled neurons in RSC subregions from the different downstream targets (*n* = 4).

**Figure S13.**
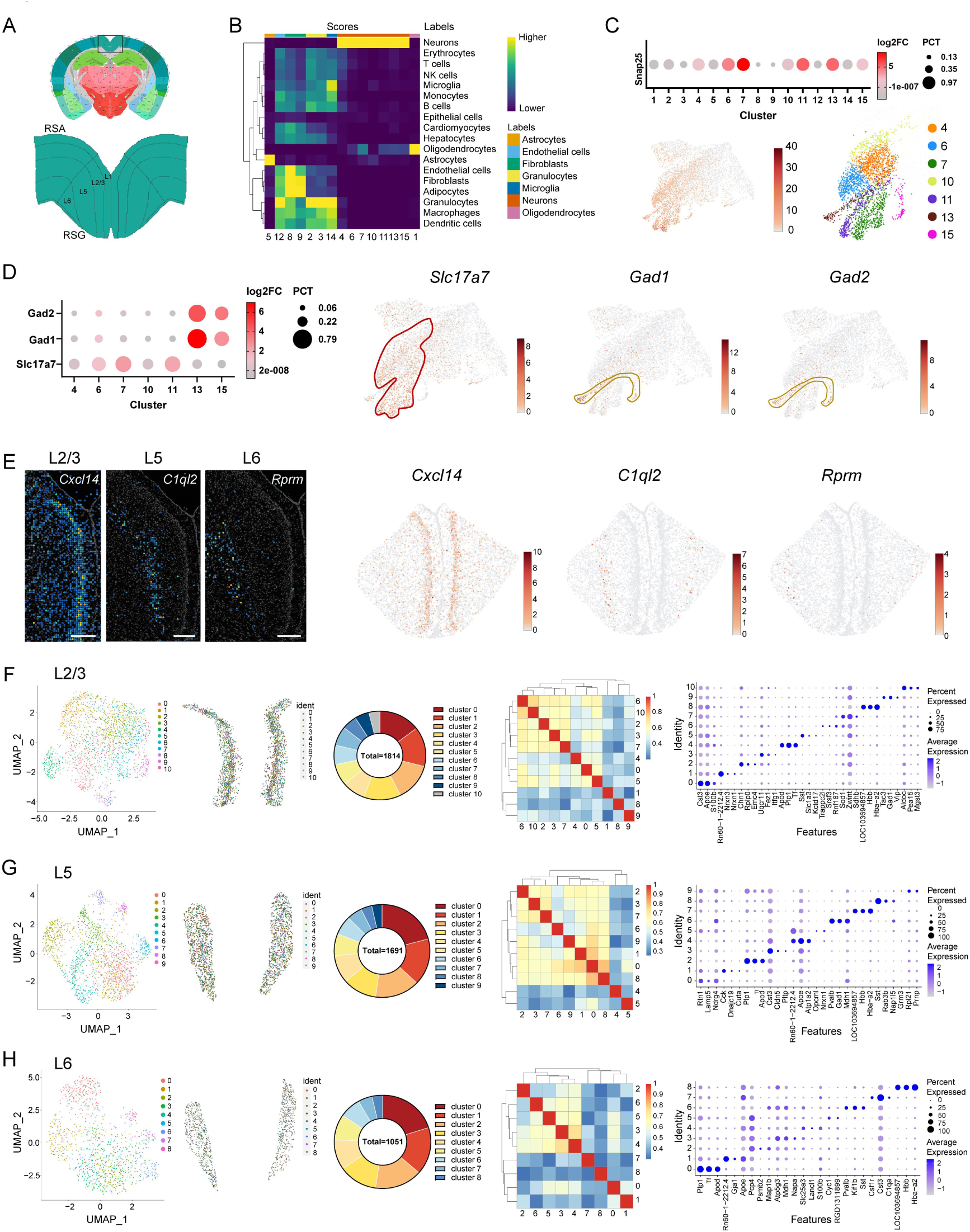
Additional information for spatial transcriptomic analysis, related to figure 5. **(A)** Layer distribution in RSG (Allen Brain Atlas). **(B)** Heatmap showing the scores defining different cell types of each cell cluster. **(C)** Top: dotplot of the average expression level and proportion of cells expressing *Snap25* in each cell cluster. Bottom: UMAP visualization of the *Snap25* expression and the identified neuronal clusters. **(D)** Left: dotplot of the average expression level and proportion of cells expressing *Gad1*, *Gad2* and *Slc17a7* in each neuronal cluster. Right: UMAP visualization of the expression of *Gad1*, *Gad2* and *Slc17a7*. **(E)** Representative images showing the expression of marker genes for layer 2/3, layer5 and layer 6 in RSG. Scale bars, 200 μm. **(F-H)** UMAP visualization, spatial distribution, proportion, co-efficiency heatmap and marker genes expression dotplot for each neuronal cluster in different layers of RSG glutamatergic neurons.

**Figure S14.**
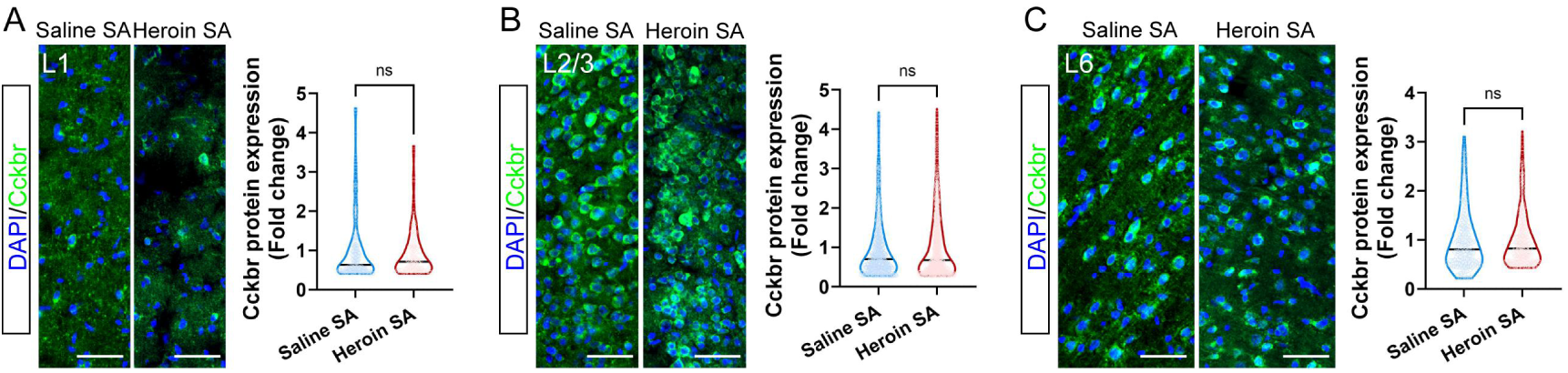
Expression of Cckbr protein across the different layers of RSG after heroin SA training, related to figure 5. **(A-C)** Left: representative images showing the expression of Cckbr protein in RSG layer 1 (A), layer 2/3 (B), layer 6 (C) of Saline SA and Heroin SA groups. Right: average expression level of Cckbr protein in RSG layer 1 (A), layer 2/3 (B), layer 6 (C) of Saline SA (*n* = 3) *vs* Heroin SA (*n* = 3) groups. Mann-Whitney test, ns, no significant difference. Scale bars, 50 μm.

**Figure S15.**
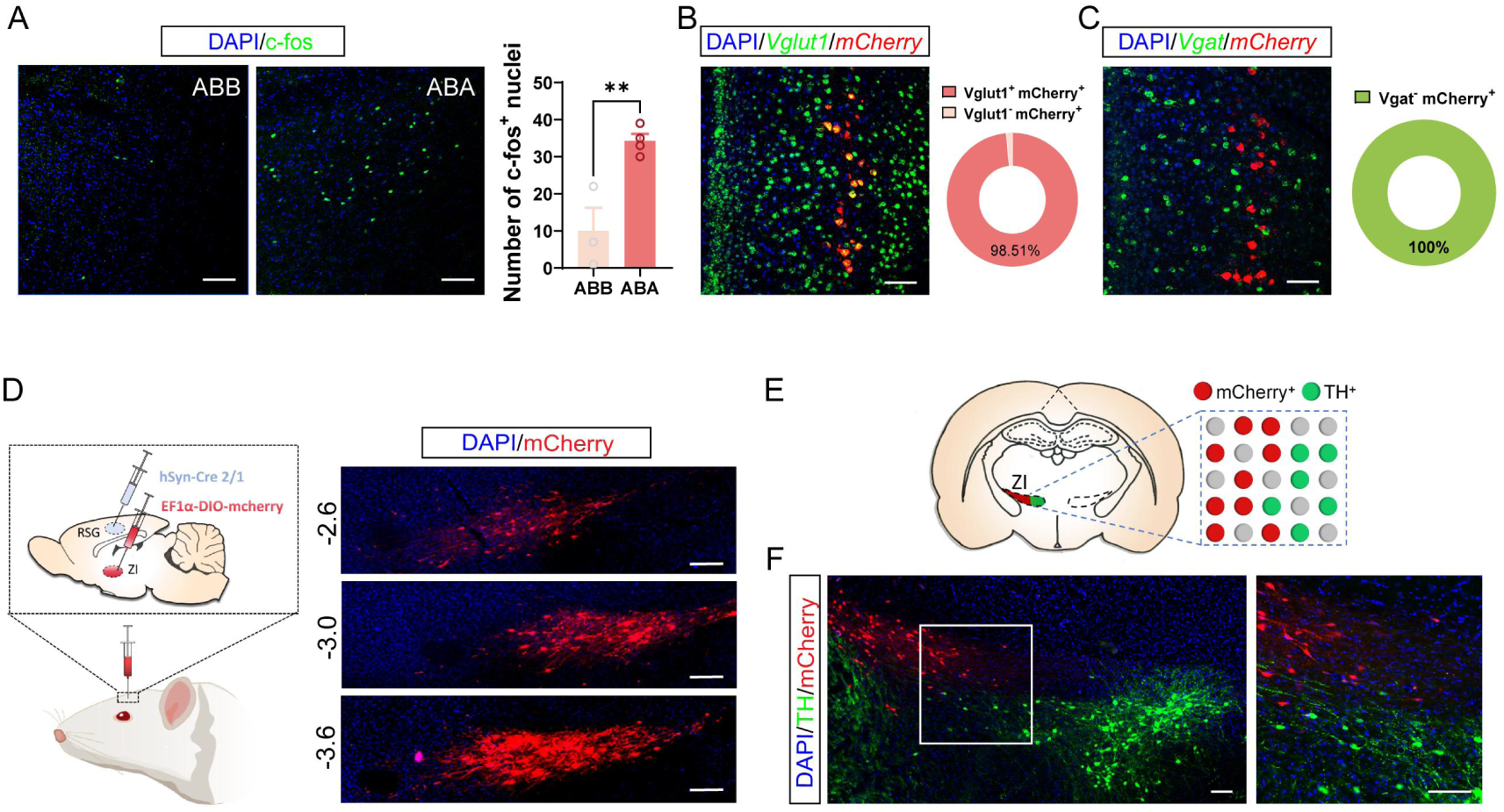
Additional information for the activation and synaptic connection of the RSG-ZI circuit, related to figure 5. **(A)** Left: representative images showing the expression of c-fos in ZI of ABB group and ABA group. Scale bars, 100 μm. Right: Number of c-fos positive nuclei in ZI of rats in ABB (*n* = 3) and ABA groups (*n* = 4). Unpaired t test, \*\**p* < 0.01. **(B)** Left: representative image showing the co-localization of retrograde-labeled *mCherry* and *Vglut1* in RSG. Scale bars, 100 μm. Right: percentage of the Vglut1-positive cells in the mCherry^+^ neurons in RSG (*n* = 3). **(C)** Left: representative image showing the co-localization of retrograde-labeled *mCherry* and *Vgat* in RSG. Scale bars, 100 μm. Right: percentage of the Vgat-negative cells in the mCherry^+^ neurons in RSG (*n* = 3). **(D)** Left: schematic showing the viral strategy for trans-synaptic anterograde tracing of RSG-ZI circuit. Right: representative images showing the RSG-innervated ZI neurons from anterior to posterior. Scale bars, 200 μm. **(E)** Schematic showing the separate distribution of mCherry-positive and dopaminergic neurons in ZI. **(F)** Representative images showing the mCherry-labeled RSG-innervated neurons and TH-positive neurons in ZI. Scale bars, 100 μm.

**Figure S16.**
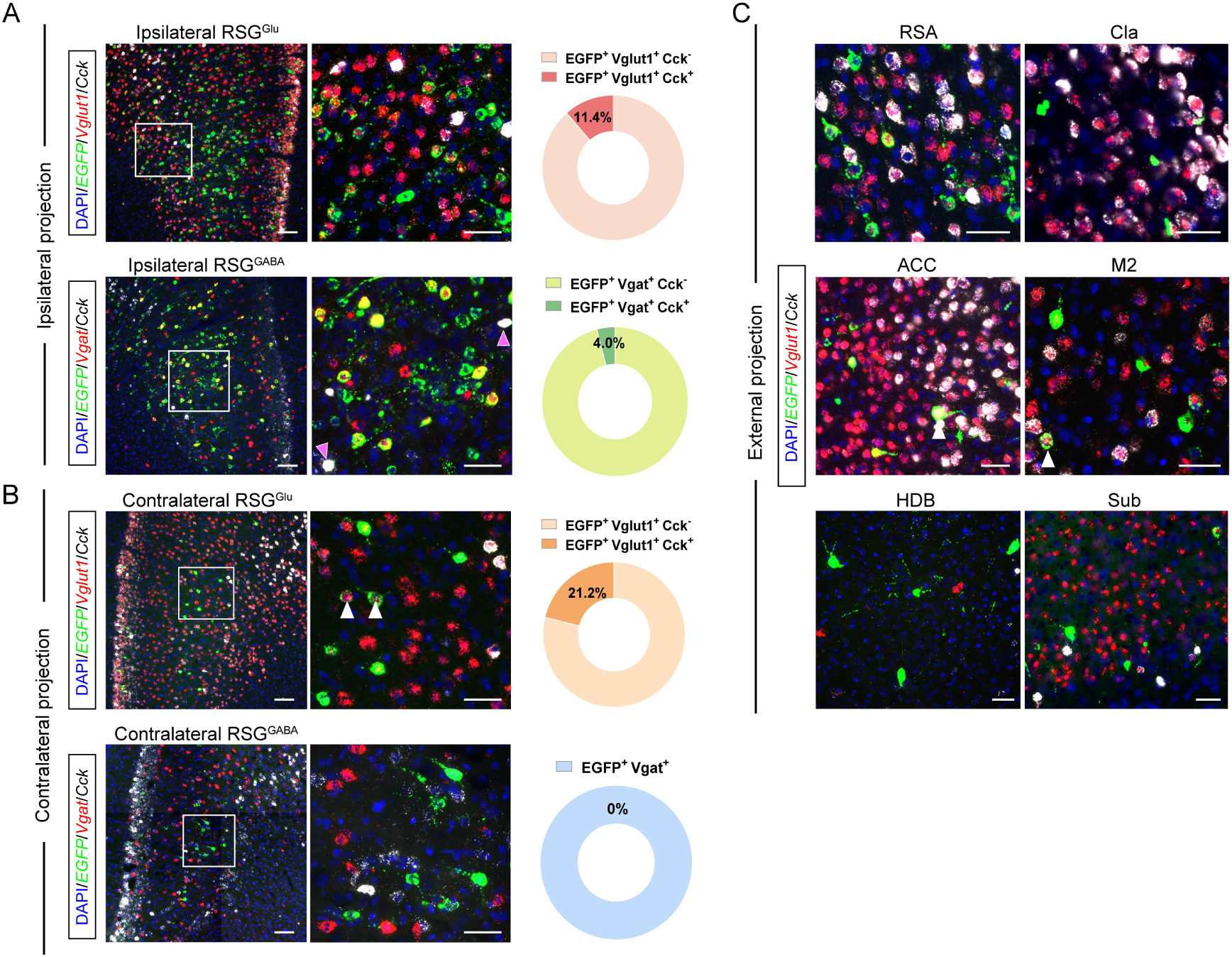
Mapping of the Cckergic input to RSG-ZI circuit, related to figure 6. **(A)** Top: representative images showing the expression of RV-labeled *EGFP*, Vglut1 and *Cck* of internal projection from ipsilateral RSG (left) and percentage of the Cck-positive cells in EGFP^+^ Vglut1^+^ neurons in RSG (right). Bottom: representative images showing the expression of RV-labeled *EGFP*, *Vgat* and *Cck* of internal projection from ipsilateral RSG (left) and percentage of the Cck-positive cells in EGFP^+^ Vgat^+^ neurons in RSG (right). Scale bars, 100 μm and 50 μm. **(B)** Top: representative images showing the expression of RV-labeled *EGFP*, *Vglut1* and *Cck* of internal projection from contralateral RSG (left) and percentage of the Cck-positive cells in EGFP^+^ Vglut1^+^ neurons in RSG (right). Bottom: representative images showing the expression of RV-labeled *EGFP*, *Vgat* and *Cck* of internal projection from contralateral RSG (left) and percentage of the Cck-positive cells in EGFP^+^ Vgat^+^ neurons in RSG (right). Scale bars, 100 μm and 50 μm. **(C)** Representative images showing the expression of RV-labeled *EGFP*, *Vglut1* and *Cck* of external projection from RSA, Cla, ACC, M2, HDB and Sub. Scale bars, 50 μm.

